# Excitatory Deep Brain Stimulation Quenches Parkinsonian Signs in the Basal Ganglia and Thalamus

**DOI:** 10.1101/2020.12.16.421024

**Authors:** Seyed Mojtaba Alavi, Amin Mirzaei, Alireza Valizadeh, Reza Ebrahimpour

## Abstract

Parkinson’s disease (PD) is associated with abnormal *β* band oscillations (13-30 Hz) in the cortico-basal ganglia circuits. Abnormally increased striato-pallidal inhibition and strengthening the synaptic coupling between subthalamic nucleus (STN) and globus pallidus externa (GPe), due to the loss of dopamine, are considered as the potential sources of *β* oscillations in the basal ganglia. Deep brain stimulation (DBS) of the basal ganglia subregions is known as a way to reduce the pathological *β* oscillations and motor deficits related to PD. Despite the success of the DBS, its underlying mechanism is poorly understood and, there is controversy about the inhibitory or excitatory role of the DBS in the literature. Here, we utilized a computational network model of basal ganglia which consists of STN, GPe, globus pallidus interna (GPi), and thalamic neuronal population. This model can reproduce healthy and pathological *β* oscillations similar to what has been observed in experimental studies. Using this model, we investigated the effect of DBS to understand whether its effect is excitatory or inhibitory. Our results show that the excitatory DBS (EDBS) is able to quench the pathological synchrony and *β* oscillations, while, applying inhibitory DBS (IDBS) failed to quench the PD signs. In light of simulation results, we conclude that the effect of the DBS on its target is excitatory.

## Introduction

Parkinson’s disease (PD) results from malfunctioning of basal ganglia (BG)^1–3^. This malfunctioning follows degeneration of dopaminergic neurons in pars compacta section of the substantia nigra (SNc)^2, 4^. Rigidity, bradykinesia, tremor, and postural instability are common signs of PD^5, 6^. In addition, this disorder is associated with excessive synchronization and abnormally *β* band (13-30 Hz) oscillations in subregions of BG^6–10^ and enhanced *β* oscillations are known as the biomarker of the PD^11–13^. However, the source of the oscillations is still under debate. Several studies have suggested the subthalamo-pallidal circuit as the main source of the generation of the *β* oscillations in experimental and computational studies^7, 8, 14–24^. While, the induction of the *β* oscillations from cortex to the BG has also been claimed^6, 25, 26^.

Furthermore, deep brain stimulation (DBS) of the BG subregions, mainly the subthalamic nucleus (STN), is a standard approach to treating PD^27–33^. Although the DBS quenches the *β* oscillations and improves the PD motor symptoms, its underlying mechanism is poorly understood^34–37^ and, there is a controversy between the excitatory and inhibitory role of the DBS.

Reduction of firing rate of the stimulated neuronal area has been observed in human^38, 39^ and monkeys^40^ with PD which remarks the inhibitory role of the DBS. Moreover, several mechanisms suggested to explain the inhibitory role of the DBS such as depolarization block^36, 41^, inactivation of voltage-gated currents^42–44^, and activation of inhibitory afferents^38, 40, 45–50^. An experimental study has suggested that the effect of the DBS is inhibitory^45^. The authors showed the GPi neurons were inhibited during stimulation, while the neurons were inhibited using GABA blocker. On the other hand, the excitatory role of the DBS has also been suggested by several studies^48, 51^. Applying DBS on the internal segment of globus pallidus (GPi) reduces firing rates of thalamic neurons which are inhibited by the GPi^52^. Also, applying the DBS on STN neurons (the excitatory neuronal population) increases the firing rate of GPi, globus pallidus externa (GPe), and substantia nigra pars reticulata (SNr) of human and animal with PD^53–55^ which support the excitatory effect of the DBS. In fact, due to the electrical pulse artifacts, the electrophysiologically investigation of an area during long term high frequency stimulation is impossible. Hence, the computational models are helpful for studying the underlying mechanism of DBS.

Computational studies have also explored the effect(s) of the DBS with inhibitory and excitatory pulses on the models of BG with PD signs. In^56^ the BG has been modelled by leaky integrate and fire (LIF) neurons which can generate *β* oscillations in PD condition, and have shown that the PD *β* oscillations quenched when the model exposed to the inhibitory DBS. While, other computational studies which are based on Hodgkin-Huxley type neurons, used excitatory DBS to suppress PD like oscillations^18, 57–62^. These studies is not shown the *β* oscillations in neuronal populations when their model is brought to PD condition. To investigate whether the effect of the DBS on its target is inhibitory or excitatory, in the current study, we used a computational model based on a variation of the model proposed in^18^ that resulted in *β* oscillations in neuronal populations in PD condition. In this study, we assumed that the subthalamo-pallidal circuit is the source of the generation of the pathological *β* oscillations. The aim of this study is to investigate whether the role of the DBS on its target is excitatory or inhibitory. Moreover, we investigate how the DBS quenches the abnormal *β* oscillations related to PD in detail. To this end we first explained the mechanisms of the *β* rhythm generation and, we found that the excitatory DBS can quench the PD signs and the inhibitory DBS fails to do so.

In this study, we mainly performed the analysis of *β* oscillations with oscillation index and also the neuronal synchronization with Fano factor in previous study^56^. Since the DBS in that study was inhibitory (the apposite results with this study), these analyses are suitable for comparison. Besides, we explored the response of thalamic neurons to cortical input that represents the thalamic fidelity which has investigated experimentally in^63^ and computationally in^64^ to more confirmation of our results.

Despite the *β* rhythm, the resting state tremor is another sign of PD that directly related to thalamic activity^65, 66^. Hence, we studied the thalamic activity in the model during the PD condition and the mechanisms of its generation. Our network model showed the excessive thalamic activity during resting state. We guess this activity is related to the Tremor. As the DBS has a therapeutic effect on resting-state tremor^35^, we also investigated the effects of DBS in our network model. We found that the excitatory DBS can reduce the excessive thalamic activity related to resting-state tremor while the inhibitory DBS cannot. In addition, we studied how the DBS can affect tremor.

## Materials and methods

### Structure of the network model

The network model consists of STN, GPe, GPi, and thalamus. Each neuronal population includes 20 Hodgkin-Huxley type neurons. The basic network model structure is similar to^18^ and^17^. the network model structure resembles the sparse pattern of connectivity^67^ proposed in^17^. STN excites GPe and GPi while receiving inhibitory input from GPe. Similar to^18^ each STN neuron receives inhibitory input from two GPe neurons. In^18^ each GPe neuron was receiving excitatory input from three STN neurons, while in our simulation, each GPe neuron receives excitatory input from one STN neuron. Similar to^18^ each GPi neuron receives excitatory input from one STN neuron. In addition, each GPi neuron receives inhibitory input from two GPe neurons. In our simulation, each thalamic neuron receives inhibitory input from one GPi neuron, while in^18^ each thalamic neuron was receiving inhibitory input from eight GPi neurons. See Figure 1 for more details on the network connectivity and structure.

**Figure 1.**
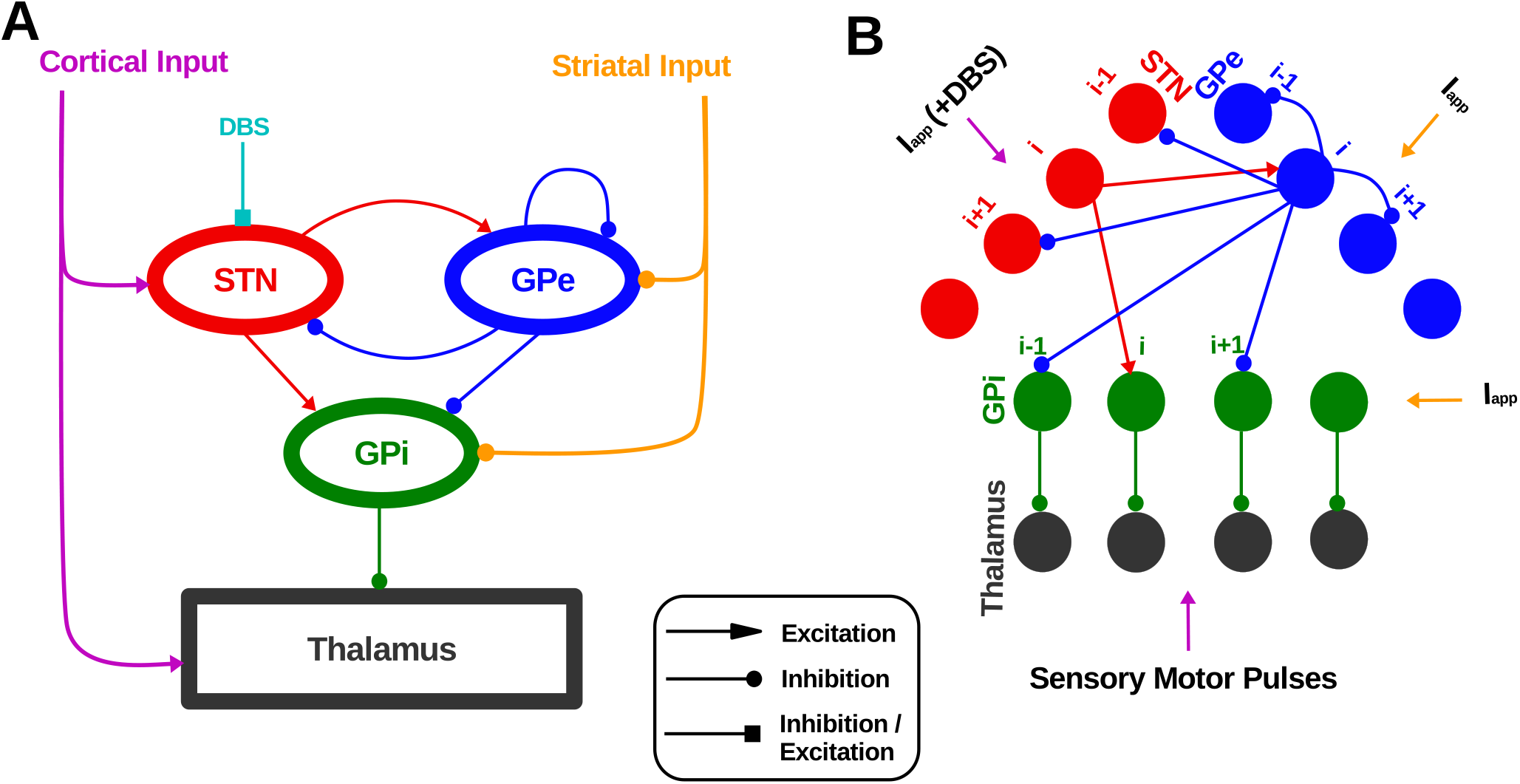
Network model structure and finding appropriate DBS currents. (A) Schematic of the network model. The DBS is considered to be either inhibitory or excitatory input to the STN. (B) Details of the network connectivity. The *i*th STN neuron excites the *i*th GPe and GPi neurons. The *i*th GPe neuron inhibits the (*i* − 1)th and (*i* + 1)th STN, GPi, and Gpe neurons. Each GPi neuron inhibits its corresponding thalamic neuron.

### Neuron and synapse model

The membrane potential of the STN, GPe, and GPi neurons in the network model was computed using the following differential equations:

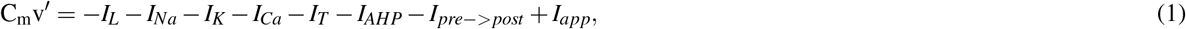

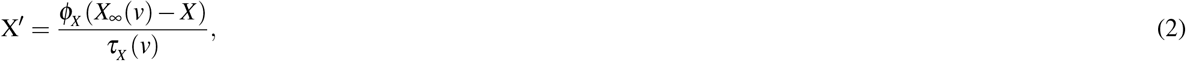

where *I*_*L*_, *I*_*Na*_, *I*_*K*_, *I*_*Ca*_, *I*_*T*_, and *I*_*AHP*_ are the leak, sodium, potassium, high threshold calcium, low threshold calcium, and after hyper polarization currents, respectively. *I*_*app*_ is the external current applied to the neurons (i.e., the DBS current). *I*_*pre*−>*post*_ is synaptic current from the presynaptic to the postsynaptic neuron. *X* represents gating channels such as potassium channels (n), opening (m) and closing (h) sodium channels, and low threshold calcium channels (r). The *τ*_*X*_ (*v*) in equation 2 is defined as follows:

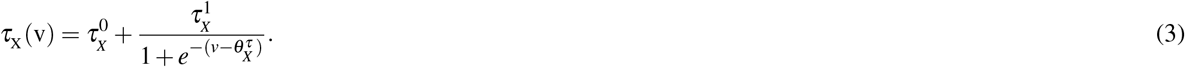

While, in the GPe and GPi neurons the *τ*_*X*_ (*v*) is constant and equal to *τ*_*r*_. The ionic currents used in equation 1 were computed as follows:

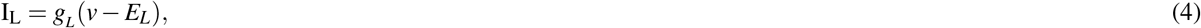

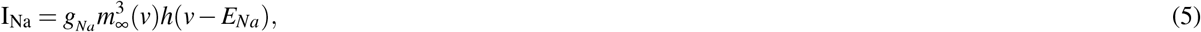

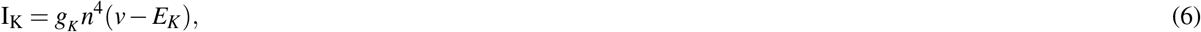

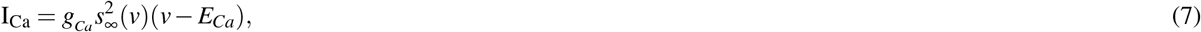

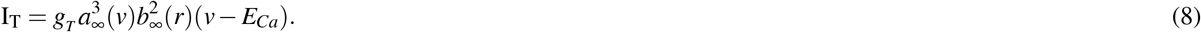

In the equations 4 to 8 the *X* = *n, h* is the ionic gating channel variables (h for closing sodium channel and n for potassium). In these equations, the *X*_∞_ = *m, a, r* or *s* is the steady-state of the ionic gating channels (m for opening sodium channel, a for T-type and s for L-type calcium channel) and is computed by the equation 9.

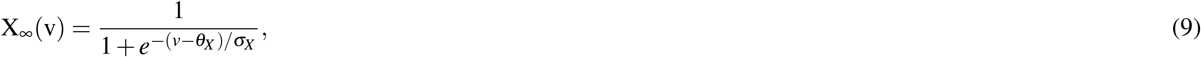

But, the function *b*_∞_(*r*) used in 8 is computed with different equation:

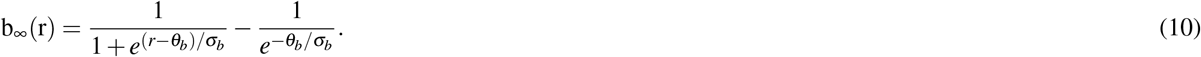

The after hyper-polarization (AHP) current used in equation 1 (*I*_*AHP*_) is

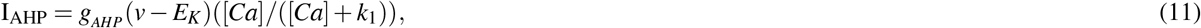

where the [*Ca*] is the intra-cellular calcium concentration:

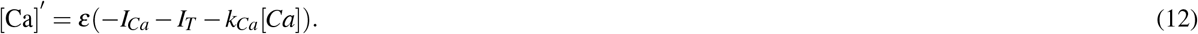

The parameters and their values of STN, GPe, and GPi neurons are presented in the table 1 to 3.

**Table 1.** Parameters and their corresponding values of STN neurons. The stars indicate transition from healthy to PD.

| Parameter | Value | Parameter | Value | Parameter | Value | Parameter | Value |
| --- | --- | --- | --- | --- | --- | --- | --- |
| $g_L$ | $2.25 \text{ nS}/\mu\text{m}^2$ | $\tau_h^0$ | 1 ms | $\theta_a$ | -63 | $\sigma_n$ | 8 |
| $g_{Na}$ | $30 \text{ nS}/\mu\text{m}^2$ | $\tau_n^0$ | 1 ms | $\theta_b$ | 0.4 | $\sigma_r$ | -2 |
| $g_K$ | $40 \text{ nS}/\mu\text{m}^2$ | $\tau_r^0$ | 40 ms | $\theta_s$ | -39 | $\sigma_a$ | 7.8 |
| $g_T$ | $0.5 \text{ nS}/\mu\text{m}^2$ | $\phi_h$ | 5 | $\theta_h^\tau$ | -57 | $\sigma_b$ | -0.1 |
| $g_{Ca}$ | $0.5 \text{ nS}/\mu\text{m}^2$ | $\phi_n$ | 5 | $\theta_n^\tau$ | -80 | $\sigma_s$ | 8 |
| $g_{AHP}$ | $9 \text{ nS}/\mu\text{m}^2$ | $\phi_r$ | 2 | $\theta_r^\tau$ | 68 | $\sigma_h^\tau$ | -3 |
| $E_L$ | -60 mV | $k_1$ | 15 | $\theta^H$ | -39 | $\theta_X \tau_n$ | -26 |
| $E_{Na}$ | 55 mV | $k_{Ca}$ | 22.5 | $\theta$ | 20 | $\sigma_r^\tau$ | -2.2 |
| $E_K$ | -80 mV | $\varepsilon$ | $3 \times 10^{-5} \text{ ms}^{-1}$ | $\alpha$ | $2 \text{ ms}^{-1}$ | $\sigma^H$ | 8 |
| $E_{Ca}$ | 140 mV | $\theta_m$ | -30 | $g_{GPe \rightarrow STN}$ | $2.2 \mapsto 7^* \text{ nS}/\mu\text{m}^2$ | $\beta$ | $0.08 \text{ ms}^{-1}$ |
| $\tau_h^1$ | 500 ms | $\theta_h$ | -39 | $E_{GPe \rightarrow STN}$ | -85 mV | $I_{app}$ | $8.4 \mapsto 3^* \text{ pA}/\mu\text{m}^2$ |
| $\tau_h^1$ | 100 ms | $\theta_n$ | -32 | $\sigma_m$ | 15 | $C_m$ | 1 |
| $\theta_r^1$ | 17.5 | $\theta_r$ | -67 | $\sigma_h$ | -3.1 | | |

**Table 2.** Parameters and their corresponding values of GPe neurons. The stars indicate transition from healthy to PD.

| Parameter | Value | Parameter | Value | Parameter | Value | Parameter | Value |
| --- | --- | --- | --- | --- | --- | --- | --- |
| $g_L$ | $0.1 \text{ nS}/\mu\text{m}^2$ | $\tau_n^0$ | 0.05 ms | $\theta_s$ | -35 | $\sigma_r$ | -2 |
| $g_{Na}$ | $120 \text{ nS}/\mu\text{m}^2$ | $\tau_r$ | 30 ms | $\theta_h^\tau$ | -40 | $\sigma_a$ | 2 |
| $g_K$ | $30 \text{ nS}/\mu\text{m}^2$ | $\phi_h$ | 0.135 | $\theta_n^\tau$ | -40 | $\sigma_s$ | 2 |
| $g_T$ | $0.5 \text{ nS}/\mu\text{m}^2$ | $\phi_n$ | 0.165 | $\theta^H$ | -57 | $\sigma_h^\tau$ | -12 |
| $g_{Ca}$ | $0.15 \text{ nS}/\mu\text{m}^2$ | $\phi_r$ | 1 | $\theta$ | 30 | $\sigma_n^\tau$ | -12 |
| $g_{AHP}$ | $30 \text{ nS}/\mu\text{m}^2$ | $k_1$ | 30 | $\alpha$ | $5 \text{ ms}^{-1}$ | $\sigma^H$ | 2 |
| $E_L$ | -55 mV | $k_{Ca}$ | 2.4 | $g_{STN \rightarrow GPe}$ | $0.01 \mapsto 0.55^* \text{ nS}/\mu\text{m}^2$ | $\beta$ | $0.14 \text{ ms}^{-1}$ |
| $E_{Na}$ | 55 mV | $\varepsilon$ | $0.0055 \text{ ms}^{-1}$ | $E_{STN \rightarrow GPe}$ | 0 mV | $I_{app}$ | $5.9 \mapsto 0.5^* \text{ pA}/\mu\text{m}^2$ |
| $E_K$ | -80 mV | $\theta_m$ | -37 | $g_{GPe \rightarrow GPe}$ | $0.01 \mapsto 0.9^* \text{ nS}/\mu\text{m}^2$ | $C_m$ | 1 |
| $E_{Ca}$ | 120 mV | $\theta_h$ | -58 | $E_{GPe \rightarrow GPe}$ | -100 mV | | |
| $\tau_h^1$ | 0.27 ms | $\theta_n$ | -50 | $\sigma_m$ | 10 | | |
| $\tau_n^1$ | 0.27 ms | $\theta_r$ | -70 | $\sigma_h$ | -12 | | |
| $\theta_h^0$ | 0.05 | $\theta_a$ | -57 | $\sigma_n$ | 14 | | |

**Table 3.** Parameters and their corresponding values of GPi neurons. The stars indicate transition from healthy to PD.

| Parameter | Value | Parameter | Value | Parameter | Value | Parameter | Value |
| --- | --- | --- | --- | --- | --- | --- | --- |
| $g_L$ | $0.1 \text{ nS}/\mu\text{m}^2$ | $\tau_n^0$ | 0.05 ms | $\theta_s$ | -35 | $\sigma_r$ | -2 |
| $g_{Na}$ | $120 \text{ nS}/\mu\text{m}^2$ | $\tau_r$ | 30 ms | $\theta_h^\tau$ | -40 | $\sigma_a$ | 2 |
| $g_K$ | $30 \text{ nS}/\mu\text{m}^2$ | $\phi_h$ | 0.1 | $\theta_n^\tau$ | -40 | $\sigma_s$ | 2 |
| $g_T$ | $0.5 \text{ nS}/\mu\text{m}^2$ | $\phi_n$ | 0.135 | $\theta^H$ | -57 | $\sigma_h^\tau$ | -12 |
| $g_{Ca}$ | $0.15 \text{ nS}/\mu\text{m}^2$ | $\phi_r$ | 1 | $\theta$ | 30 | $\sigma_n^\tau$ | -12 |
| $g_{AHP}$ | $30 \text{ nS}/\mu\text{m}^2$ | $k_1$ | 30 | $\alpha$ | $5 \text{ ms}^{-1}$ | $\sigma^H$ | 2 |
| $E_L$ | -55 mV | $k_{Ca}$ | 2.4 | $g_{STN \rightarrow GPe}$ | $0.005 \mapsto 1.1^* \text{ nS}/\mu\text{m}^2$ | $\beta$ | $0.14 \text{ ms}^{-1}$ |
| $E_{Na}$ | 55 mV | $\varepsilon$ | $0.0055 \text{ ms}^{-1}$ | $E_{STN \rightarrow GPe}$ | 0 mV | $I_{app}$ | $7.7 \mapsto 4^* \text{ pA}/\mu\text{m}^2$ |
| $E_K$ | -80 mV | $\theta_m$ | -37 | $g_{GPe \rightarrow GPe}$ | $0.01 \mapsto 1.9^* \text{ nS}/\mu\text{m}^2$ | $C_m$ | 1 |
| $E_{Ca}$ | 120 mV | $\theta_h$ | -58 | $E_{GPe \rightarrow GPe}$ | -100 mV | | |
| $\tau_h^1$ | 0.27 ms | $\theta_n$ | -50 | $\sigma_m$ | 10 | | |
| $\tau_n^1$ | 0.27 ms | $\theta_r$ | -70 | $\sigma_h$ | -12 | | |
| $\theta_h^0$ | 0.05 | $\theta_a$ | -57 | $\sigma_n$ | 14 | | |

The membrane potential of thalamic neurons in the network model is computed using the following differential equations:

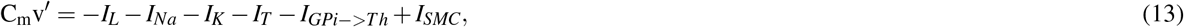

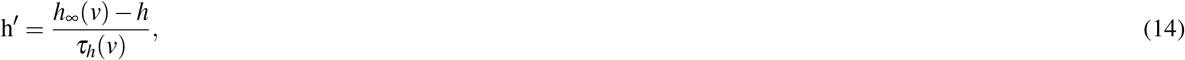

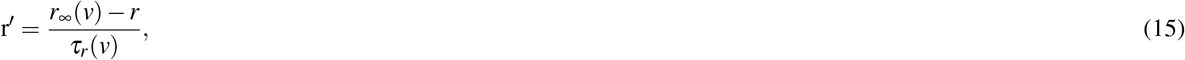

where *I*_*L*_, *I*_*Na*_, *I*_*K*_, and *I*_*T*_ are the leak, sodium, potassium, and low threshold calcium currents, respectively. *I*_*GPi*−>*Th*_ is the synaptic current from a GPi neuron to a thalamic neuron in the network model. The *I*_*SMC*_ represents cortico-thalamic sensorimotor pulses applied to the thalamic neurons. The equations 16 to 19 compute the ionic currents used in equation 13.

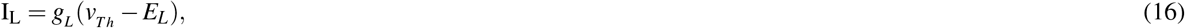

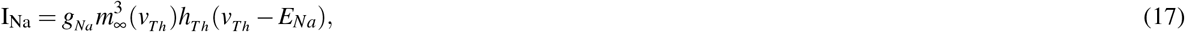

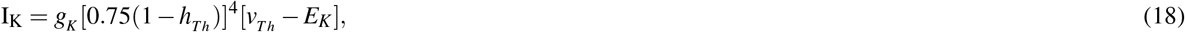

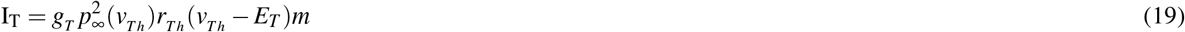

and the functions used in equations 14 to 19 are computed as follows:

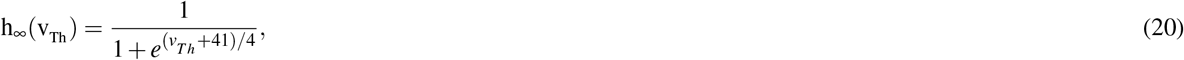

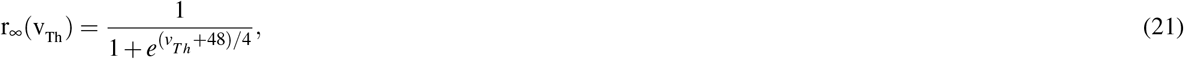

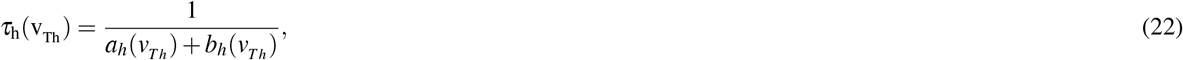

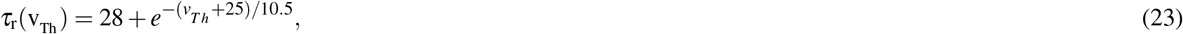

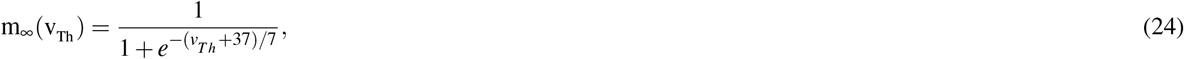

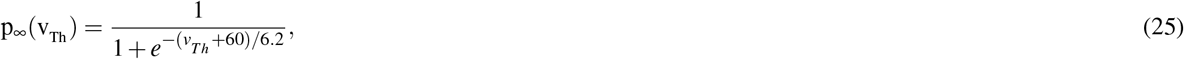

which the *a*_*h*_(*v*_*Th*_) and *b*_*h*_(*v*_*Th*_) are

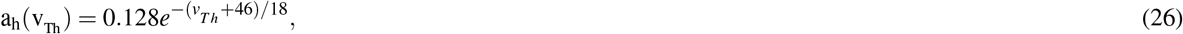

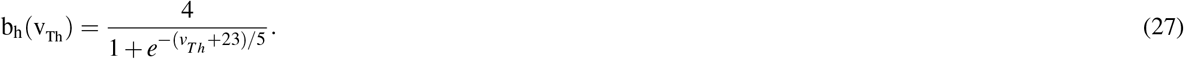

The parameters and their values of thalamic neurons are presented in table 4.

**Table 4.** thalamic parameters and the corresponding values

| Parameter | Value |
| --- | --- |
| $g_L$ | $0.05 \text{ nS}/\mu\text{m}^2$ |
| $E_L$ | -70 mV |
| $g_{Na}$ | $3 \text{ nS}/\mu\text{m}^2$ |
| $E_{Na}$ | 50 mV |
| $g_T$ | $5 \text{ nS}/\mu\text{m}^2$ |
| $E_T$ | 0 mV |
| $g_{GPi \rightarrow Th}$ | $0.005 \text{ nS}/\mu\text{m}^2$ |
| $E_{GPi \rightarrow Th}$ | -85 mV |
| $\theta^H$ | -57 |
| $\sigma^H$ | 2 |

In addition to modification of the network structure we modified the network model parameters compared to the Terman et al. (2004) as follows: for the STN neurons *g*_*Na*_ was decreased from 37.5 to 30 *nS/µm*^2^. The *g*_*K*_ was decreased from 45 to 40 *nS/µm*^2^. The value of *ϕ* was taken from^68^ (i.e., *ϕ*_*n*_ = *ϕ*_*h*_ = 5, *ϕ*_*r*_ = 2). The value of *ε* was considered to be 3 × 10^−5^ *ms*^−1^. The *g*_*GPe* − >*STN*_ and *I*_*app*_ of the STN in the healthy state is 2.2 *nS/µm*^2^ and 8.4 *pA/µm*^2^, respectively. In the PD state of the network model these two parameters were changed to 7 *nS/µm*^269–72^ and 3 *pA/µm*^273^, respectively.

To simulate GPe neurons in the network model, we set *ϕ*_*h*_ = 0.135, *ϕ*_*n*_ = 0.165, *ϕ*_*r*_ = 1, and *ε* = 0.0055 (similar to^68^). The *g*_*GPe*−>*GPe*_, *g* _*STN*−>*Gpe*_, and *I*_*app*_ of GPe in the healthy state were 0.01, 0.01 *nS/µm*^2^, and 5.9 *pA/µm*^2^, respectively. To simulate the PD state of the network model, these parameters were changed to 0.9, 0.55*nS/µm*^269–72^, and 0.5 *pA/µm*^2^, respectively. Note that in the PD state of the network model *I*_*app*_ of the GPe and GPi decreases leading to less activity of the GPe and Gpi neurons due to the increasing striatal inhibition (explained in^56^) in the network model. Parameters of the GPi neurons are similar to the GPe neurons with the difference that for the GPi neurons, *ϕ*_*h*_ = 0.1 and *ϕ*_*n*_ = 0.135. The *g*_*GPe*−>*GPi*_, *g*_*STN*−>*GPi*_, and *I*_*app*_ of GPi in the healthy state were 0.01, 0.005*nS/µm*^2^, and 7.7 *pA/µm*^2^. To simulate the PD state of the network model, these parameters similar to GPe neurons were changed to 1.9, 1.1*nS/µm*^2^, and 4 *pA/µm*^2^, respectively. Parameters of the thalamic neurons are the same as in^18^ with the difference that in our network model *g*_*GPi*−>*Th*_ was 0.05 *nS/µm*^2^. These modifications moved our network model activity more close to the experimental results.

The synaptic model used here is of a conductance-based type similar to the model used in^17, 18, 61, 62, 68, 74^. The synaptic currents used in equations 1 and 13 are computed as follows:

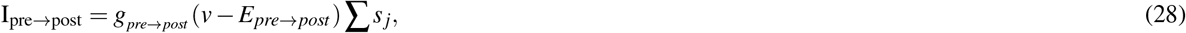

where the *j* is the index of the presynaptic neuron. The parameter *s* in equation 28 is

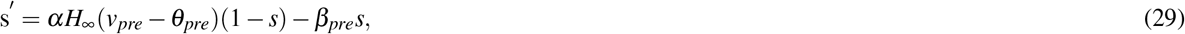

where the *H*_∞_(*v*) as follows:

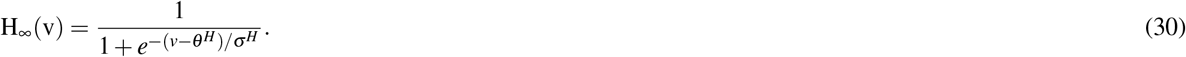

### Population Firing Rate

To compute the time resolved population firing rate for each neuronal population in the network model we used 10 milliseconds sliding window and shifted with steps of 1 millisecond over the entire simulation time while for each step we counted the number of spikes for all neurons in the population and converted it to spikes per second (i.e., to *Hz*).

### Sensorimotor and DBS pulses

The sensorimotor and DBS pulses are simulated using the following equation:

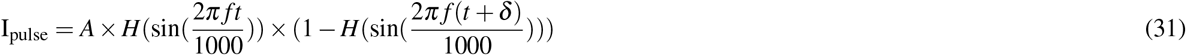

where *A, f, t*, and *δ* are pulse amplitude, frequency, time (in *ms*), and pulse duration (in *ms*), respectively. The *H*(.) is the Heaviside function. To simulate DBS, we set *f* = 150*Hz* and *δ* = 0.1*ms*. For the A parameter, we set the various values to investigate the DBS current amplitudes (see results and Figure 4D). The DBS parameters are adopted from^10, 54, 63, 75–78^. For sensorimotor pulses (i.e., cortico-thalamic input), *A* = 4.5 *pA/µm*^2^, *f* = 20*Hz*, and *δ* = 5 *ms*. To generate irregular pulses we used Poisson process in equation 31.

### Thalamic Fidelity

Thalamic neurons in the network model show four types of responses to the cortico-thalamic input pulse: 1) Correct spike: a single thalamic spike in response to a cortico-thalamic input pulse. 2) Missed spike: refers to the case when there is no thalamic spike in response to a cortico-thalamic input spike. 3) Extra spike: refers to the case when a thalamic neuron shows more than one spike in response to a cortico-thalamic input pulse. 4) Undesired spike: occurs when a thalamic neuron spikes while there is no cortico-thalamic input spike (Supplementary figure 1). According to the four response types of the thalamic neurons to a single cortical pulse, the thalamic fidelity is computed using the following equation:

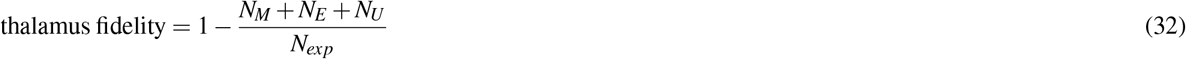

where the *N*_*M*_, *N*_*E*_, and *N*_*U*_ are the number of missed spikes, extra spikes, and undesired spikes, respectively. The *N*_*exp*_ is the number of expected thalamic spikes due to cortico-thalamic pulses. Since each cortical pulse is given to all thalamic neurons in the network model, we expect to observe that each thalamic neuron, relaying the cortico-thalamic pulse, emits a spike in response to the cortico-thalamic pulse. Therefore, the number of expected thalamic spikes in response to the cortical inputs (i.e., *N*_*exp*_) equals the number of thalamic neurons multiplied by the number of cortico-thalamic pulses^18, 79^.

### Synchrony index

We used Fano Factor (FF) to measure synchronous spiking activity for each neuronal population in the network model. To compute FF we used the following equation:

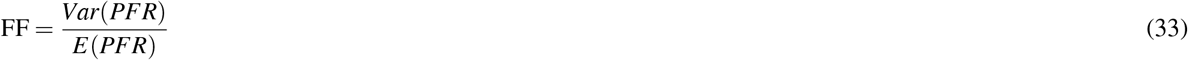

where the *Var*(.) and *E*(.) are the variance and mean of the population firing rate (*PFR*), respectively. Higher FF values represent more synchrony in the spiking activity of a neuronal population in the network model^56, 80^.

### Mean power spectral density

The power spectral density (PSD) of the population firing rates was computed using Welch’s method in python 2.7 (i.e., using scipy.signal.welch python package;^81^). The sampling rate and the segment length were set to 1000 Hz and 1000 data points, respectively. Other parameters required for the scipy.signal.welch function were set to the predefined default values (see https://docs.scipy.org/doc/scipy-0.14.0/reference/generated/scipy.signal.welch.html). The mean power spectral density was computed by averaging over 50 simulations.

### Oscillation index

The oscillation index was computed by dividing the area under the curve of a PSD in the *β* frequency range (i.e., between 13 to 30 Hz) by the area under the curve for the whole frequency range (i.e., from 1 to 500 Hz).

### Tremor frequency

We assumed that the extra spikes of the thalamic neurons during the resting state of the network model (i.e., when there are no cortico-thalamic sensorimotor pulses) are related to limb tremor. So, to compute the tremor frequency, we measured the mean firing rate of thalamic neurons (in Hz), over the whole simulation period, during the resting state.

### Simulation

The simulations were implemented in python 2.7. All differential equations were solved using odeint from SciPy library^81^ with 0.05 *ms* time resolution (see https://docs.scipy.org/doc/scipy-0.18.1/reference/generated/scipy.integrate.odeint.html). To reduce the simulation time, we performed parallel programming using the python message passing interface (MPI) in cluster computing with 30 core processors (Intel 3.2 *GHz*). To avoid the initial transient network model responses, we did not consider the first 250 *ms* of each simulation in our analysis.

## Results

### The network model captures features of the healthy and PD BG

The activity of different BG regions in the healthy state is non-oscillatory and desynchronized^3, 82–84^. This feature is captured in our network model. Similar to the experimental results, the STN spiking activity in the healthy network model is asynchronous irregular (Figure 2A, top panel; the same for the GPe and GPi spiking activity; Supplementary figure 2A and D). This is also reflected in the STN population activity in the healthy state of the network model (Figure 2B, top panel, and also Supplementary figure 2B and E, top panels). The STN population activity in the healthy network model is non-oscillatory. This leads to a flat PSD of the STN population firing rate in the healthy state (Figure 2C, Supplementary figure 2C and F). Altogether, these results indicate that the healthy activity of the network model is irregular, in line with the experimental studies showing that the BG activity in the healthy state is non-oscillatory.

**Figure 2.**
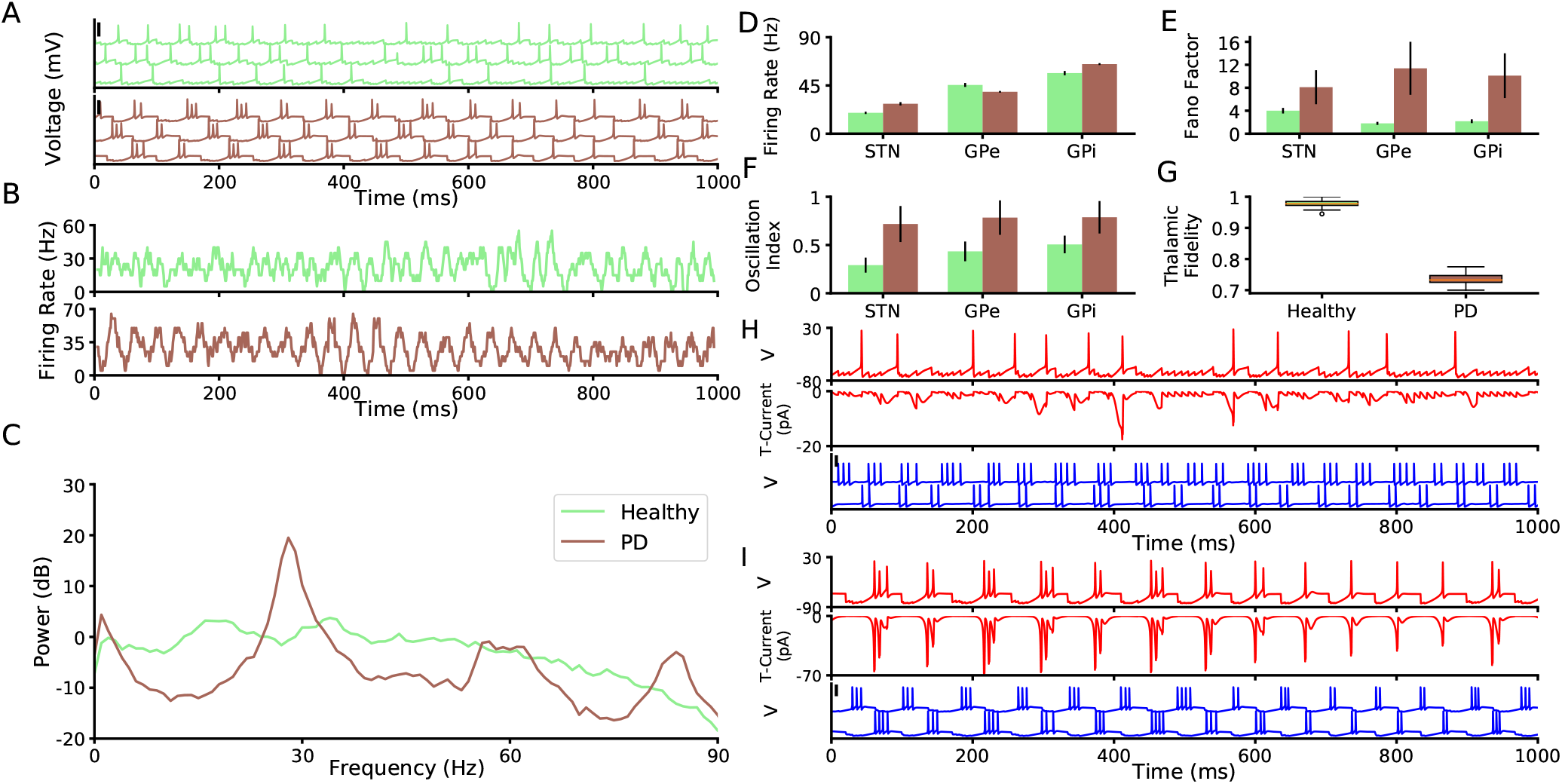
Neuronal and population properties of BG in healthy and PD states. (A) Membrane potential of three STN neurons in the network model in the healthy state (top) and PD state (bottom). The black vertical thick lines indicate 50 mV. (B) Time resolved population firing rate of the STN neurons in the Healthy state (top) and PD state (bottom). (C) Mean power spectrum (average of 50 trials) of the STN time resolved population firing rate in the healthy state (light green) and PD state (brown). (D-F) Population mean firing rate (D), Fano factor (E), and oscillation index (F) of the STN, GPe, and GPi in the network model (error bars show standard deviation; color codes correspond to C). (G) thalamic fidelity in the healthy state, PD state, and during regular and irregular EDBS. (H) The membrane potential of an STN neuron (top), its corresponding T-type calcium current (middle), and two connected GPe neurons to the STN neuron (bottom) in a healthy state. (I) The same as H for PD state.

The STN, GPe, and GPi mean population firing rates, in the healthy state of the network model, are 19.4±1.1*Hz*, 45.47±1.2*Hz* and 56.52±2*Hz*, respectively which match the previously reported experimental values^84, 85^.

In the PD state, the network model neurons show synchronized bursts of spiking activities in the *β* frequency range (Figures 2A and B, and also Supplementary figure 2A,B,D, and E, bottom panels). This also matches experimental studies which indicate synchronized *β* band oscillatory spiking activities in the BG as a hallmark of PD^7–10, 85–99^.

To bring the network model from the healthy to the PD state, we followed three steps. First, the *I*_*app*_ applied to the GPe and to the GPi neurons (Equation 1) was decreased from 5.9 *pA/µm*^2^ and 7.7 *pA/µm*^2^ (healthy state) to 0.5 *pA/µm*^2^ and 4 *pA/µm*^2^(PD state), respectively. This reduction represents the increase in the striato-pallidal inhibition^3, 19, 100^. This leads to a reduction in the activity of the GPe neurons (39.1±0.8*Hz*; independent two-tailed t-test, *p* < 0.001) in the PD state of the network model, compared to the healthy state (Figure 2D). This is in line with the experimental studies indicating that the GPe firing activity decreases during PD^84, 101, 102^. However, despite decreasing the *I*_*app*_ applied to the GPi neurons, the GPi firing rate increases (64.9±0.86*Hz*; independent two-tailed t-test, *p* < 0.001) compared to the healthy state (Figure 2D;^101, 103, 104^). The reason is that the lower GPe activity during the PD state of the network model disinhibits the GPi neurons. Thereby, the GPi population firing rate in the PD network model increases compared to the healthy state (Figure 2D).

Second, the *I*_*app*_ applied to the STN neurons, representing cortico-subthalamic input, in the network model (Equation 1) was decreased from 8.4 *pA/µm*^2^ (healthy state) to 3 *pA/µm*^2^ (PD state). Such a change in the network model is in line with the experimental studies showing that the cortical activity decreases in PD^73^ which can lead to less cortico-subthalamic drive, due to direct cortico-subthalamic connectivity^105–107^. Note that despite decreasing the *I*_*app*_ of the STN neurons in the PD state of the network model, the STN activity increases (27.9±1.5*Hz*; independent two-tailed t-test, *p* < 0.001) compared to the healthy state (Figure 2D)^85, 86^. The reason for this is STN disinhibition due to a reduction in the activity of the GPe units in the PD state of the network model (Figure 2D).

Third, the synaptic connections in the subthalamo-pallidal circuit (STN to GPe and GPe to STN synapses) were strengthened in the PD state, compared to the healthy state (see Materials and methods). Such subthalamo-pallidal synaptic strengthening is in line with the experimental studies showing that both STN to GPe and GPe to STN synapses are strengthened during PD^69–72^.

Applying these three changes brings the network model from the healthy state to the PD state. STN neurons of the PD network model show synchronized bursts of spiking activity in the *β* frequency range (Figure 2A; the GPe and GPi neurons show the same behaviour; Supplementary figure 2A and D). The STN PD-like *β* band oscillations are also observed in the STN population firing rate (Figures 2B and C) as well as in the GPe and GPi population activities (Figures 2E and F, and also see Supplementary figure 2).

The pallido-thalamic pathway is important for flowing information (i.e., the motor information) that is affected by abnormal activity in BG as what has been observed in the previous experimental study^63^. Also, the poor connectivity of the thalamus has been shown in fMRI study in rats with PD^108^. Therefore, we simulated the thalamic information processing (i.e., corticothalamic motor commands) by measuring thalamic fidelity (see Materials and method, and see also the Supplementary figure 1) in the healthy and PD states of the network model. Our results indicate that thalamic fidelity (see materials and methods) to the cortical motor commands decreases in the PD state of the network model compared to the healthy state (Figure 2G). This result is in line with the previous computational studies^18, 59, 60, 79, 109^ indicating a reduction of the thalamic fidelity during PD. This reduction comes from the increasing burst rate of the GPi neurons compared to the healthy state (See section EDEBS and IDBS Effects) that input to the thalamus. This observation matches with the previous experimental^54^ and computational studies^61^ and confirm the validity of our results.

The reason why the model exhibits the *β* oscillations in the PD state is the rebound bursting activity of the STN neurons due to receiving pallidal inhibition. In line with the experimental studies^88, 110, 111^, the STN neurons in the network model show rebound bursts after the strong inhibition. When an STN neuron receives the strong burst from a GPe neuron, its T-type calcium current increases to the sufficient value that results in rebound burst of the STN neuron (Figure 2I). The resulted STN burst goes back to the GPe neurons through the excitatory subthalamo-pallidal pathway and reverberates the bursting behaviour. This mechanism leads the *β* oscillations the population activity. While, in the healthy state, the pallido-subthalamic burst input is not high enough to elicit rebound burst acitivty in the stn neurons (Figure 2H).

All in all, our network model can reproduce features of the experimental data for both the healthy and PD state of the BG. Mainly, the network model shows asynchronous irregular spiking activity in the healthy state and synchronous *β* band oscillatory activity in the PD state. In addition, thalamic fidelity decreases in the PD state of the network model compared to the healthy state.

### Effects of EDBS and IDBS on the network model

The STN high frequency DBS has therapeutic effects on PD signs such as reducing the pathological *β* oscillations in the cortico-BG loop^33, 36, 76, 112–114^, and improving PD-related motor symptoms^27–32, 115–117^. However, the mechanism(s) underlying STN DBS is yet unknown. To understand whether the STN DBS therapeutic effects are due to excitation of STN or inhibition of STN, we applied EDBS (i.e., excitatory DBS) and IDBS (i.e., inhibitory DBS) to the STN neurons in the network model and investigated the effect(s) of each DBS type. We tested which DBS type can quench the PD-like *β* oscillations in the network model. Furthermore, we also tested which DBS type can improve the thalamic fidelity to the cortical motor commands.

To this end, we applied various current amplitudes in both types of DBS to STN to see their effect on *β* oscillations. We found that the EDBS can quench the *β* oscillations in STN and GPe populations with high enough amplitude (approximately 120 pA; Figure 3 and Supplementary figure 3A and B, top panels), while the IDBS failed to quench the *β* oscillations in all applied current amplitudes (Figure 3 and Supplementary figure 3A and B, bottom panels).

**Figure 3.**
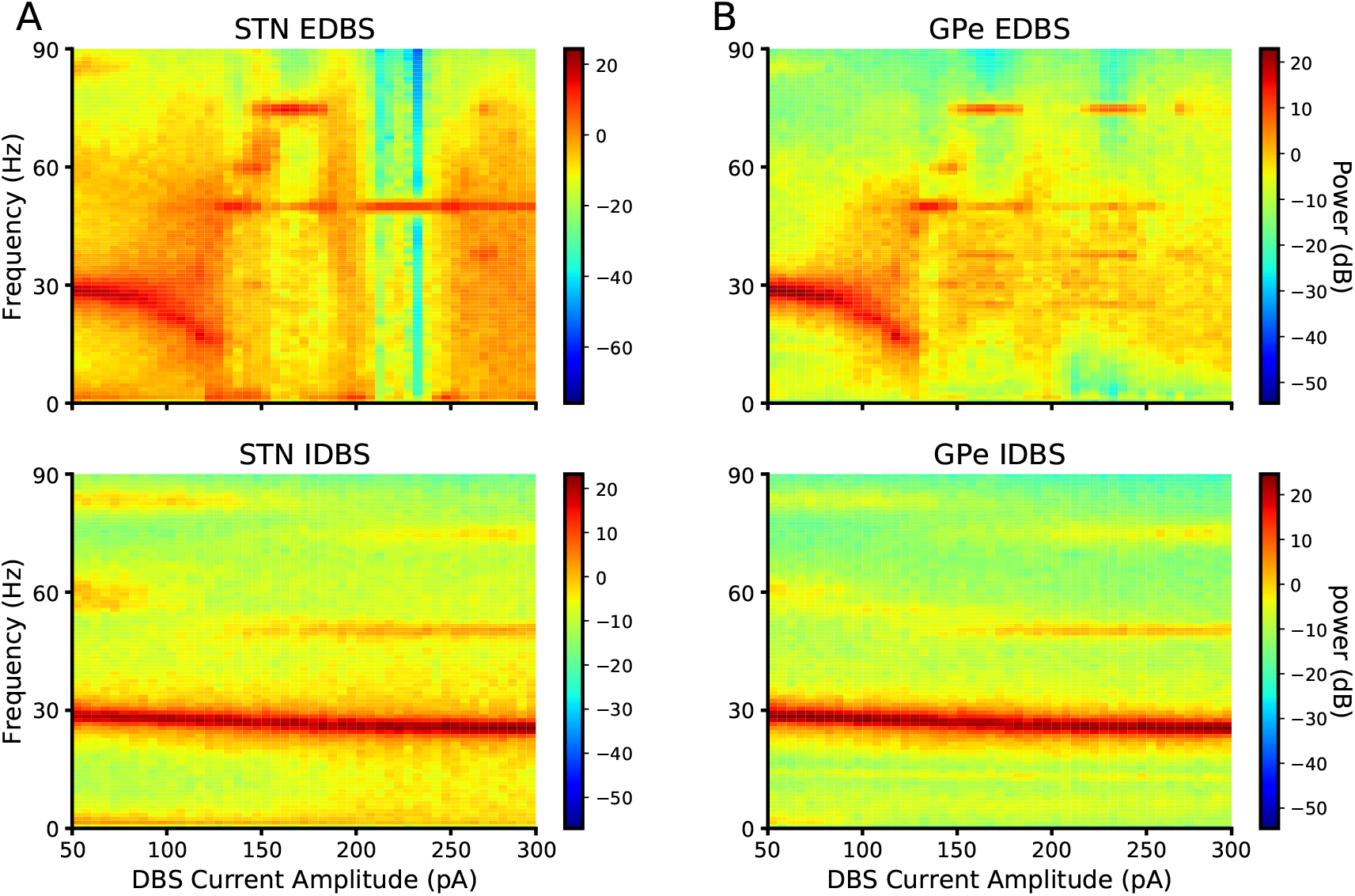
Spectrogram of STN and GPe in EDBS and IDBS states. (A) The spectrogram of STN across DBS current amplitude in EDBS (top) and IDBS (bottom) states. (B) The same as A for GPe. Each point in all spectrograms is averaged over 20 trials.

To investigate the DBS effects, we chose the specific values of EDBS and IDBS current amplitudes. For a better comparison of two DBS types, we found the amplitudes of the DBS that the STN mean firing rate to be approximately in the same values. To this end, we computed the STN mean firing rate (over 20 trials) by varying *A* = 50 *pA/µm*^2^ to 300 *pA/µm*^2^ (Figure 4-D), then we chose the constraint that the firing rate of the STN in excitatory DBS (EDBS) and inhibitory DBS (IDBS) to be equal in the cases. Therefore, to simulate EDBS, we set *A* = 126.57 *pA/µm*^2^ and 147.36 *pA/µm*^2^, *f* = 150*Hz*. Parameter settings for IDBS were the same as EDBS, with the difference that *A* = − 126.57 *pA/µm*^2^ and − 147.36 *pA/µm*^2^. Due to the similarity of results, we reported the DBSs with 147.36*pA* current except for the Figure 6.

**Figure 4.**
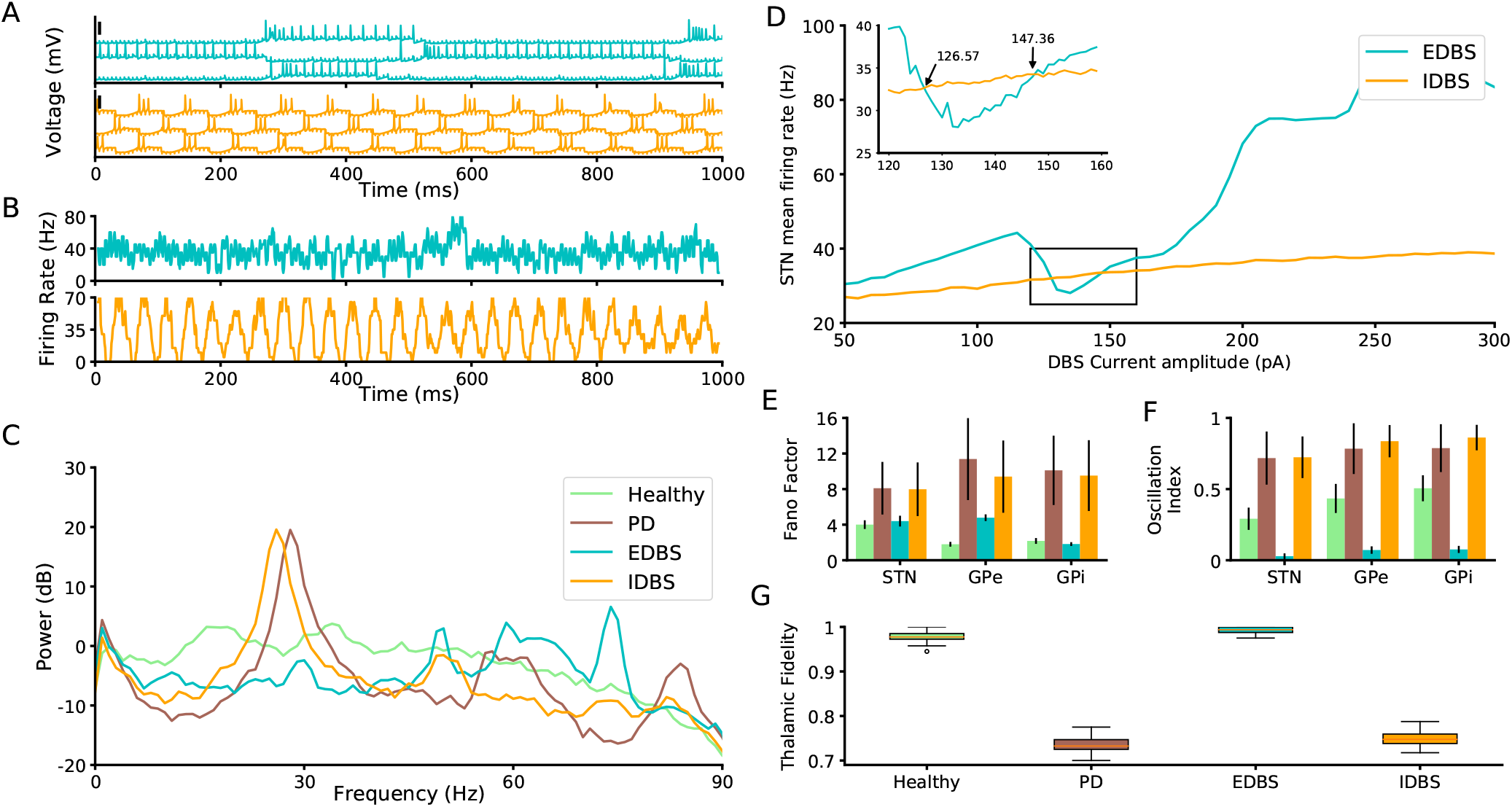
Neuronal and population properties of BG in EDBS and IDBS states. (A) Membrane potential of three STN neurons in the network model in the EDBS (top) and IDBS (bottom). The black vertical thick lines indicate 50 mV. (B) Time resolved population firing rate of the STN neurons in the EDBS (top) and IDBS (bottom). (C) Mean power spectrum (average of 50 trials) of the STN time resolved population firing rate in the healthy state (light green), PD state (brown), and during EDBS (cyan) and IDBS (orange). (D) The STN mean firing rates for IDBS and EDBS is shown versus the amplitude of the stimulation current. Inset is the zoomed-in presentation of the results in the rectangle. Each point in the plots is averaged over 20 trials. The IDBS curve crosses the EDBS curve at 126.57 and 147.36 pA points. (E and F) Fano factor (E), and oscillation index (F) of the STN, GPe, and GPi in the network model (error bars show standard deviation; color codes correspond to C).(G) thalamic fidelity in the healthy state, PD state, and during regular and irregular IDBS.

**Figure 5.**
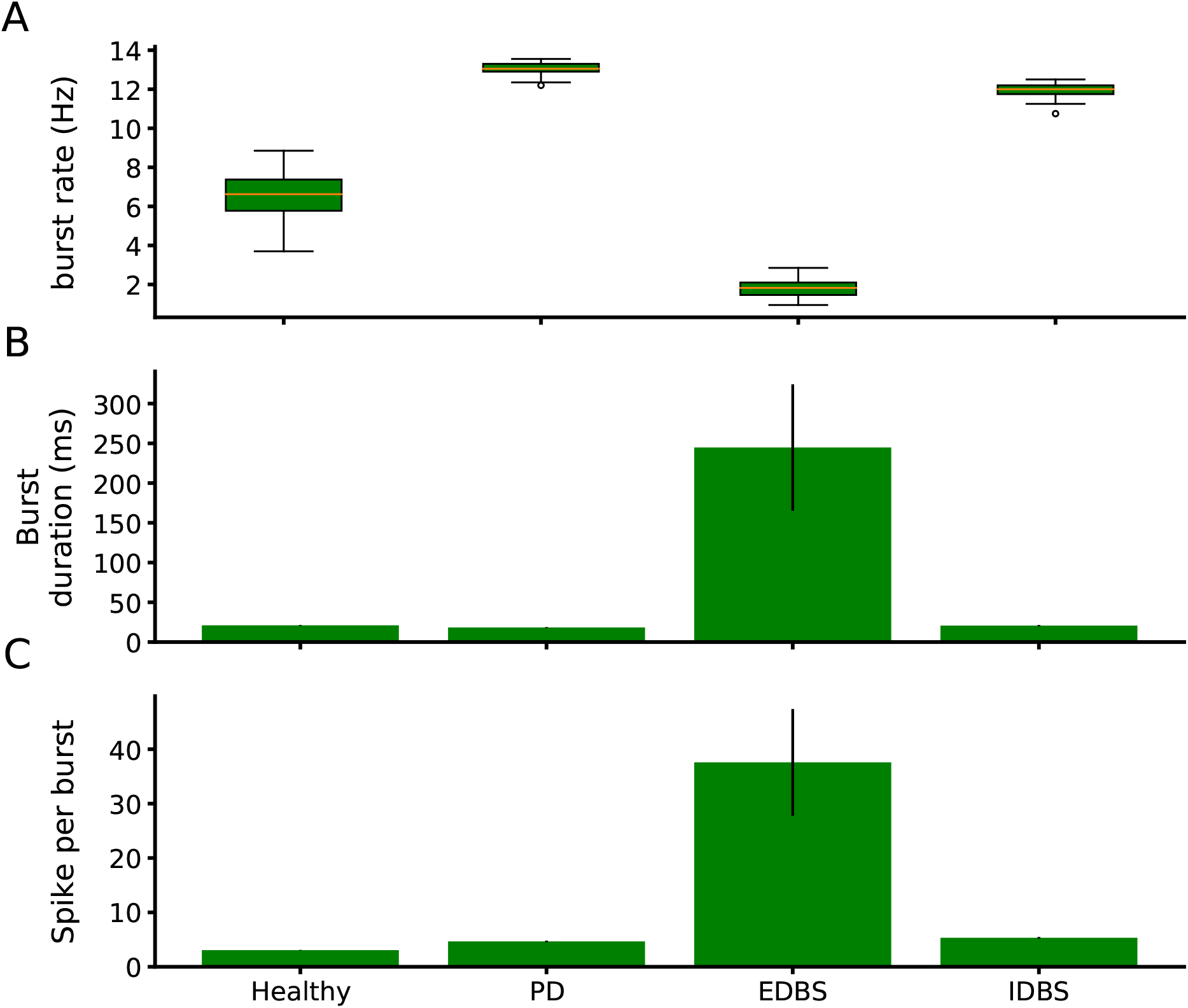
Burst profile of GPi neurons. (A) Burst rate of GPi neurons in healthy, PD, regular EDBS, irregular EDBS, regular IDBS and irregular IDBS states. (B and C) Mean burst duration (in msec) and mean the number of spikes per burst in healthy, PD, regular/irregular EDBS, and regular/irregular IDBS states. The error bars show standard deviation.

**Figure 6.**
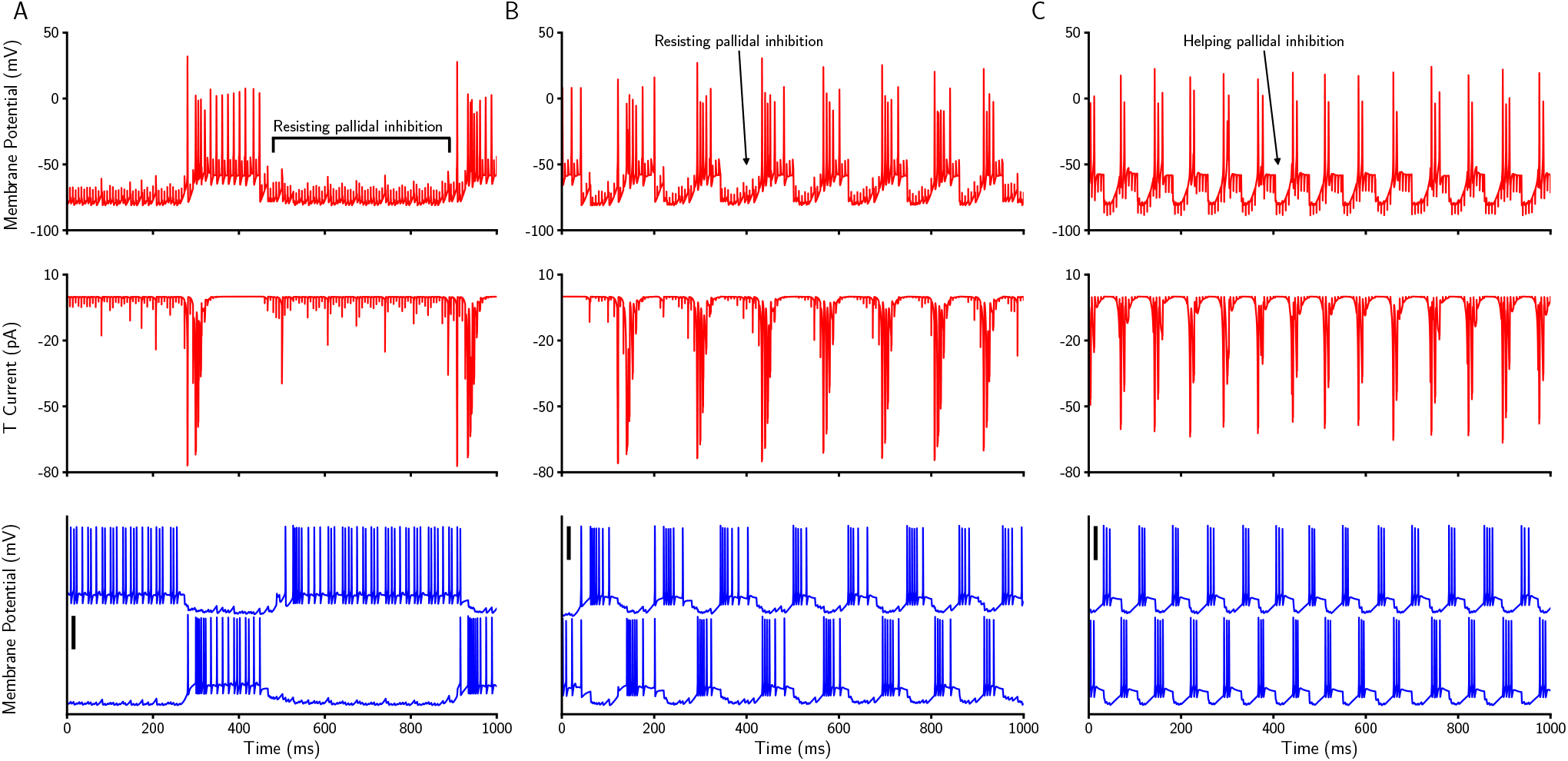
The role of inhibitory rebound bursting in STN neurons during EDBS and IDBS. (A) Membrane potential of an STN neuron (top), its corresponding T-type calcium current (middle; see materials and methods), and two connected GPe neurons to the STN neuron during EDBS with the current of 147.36*pA*. (B) The same as A, EDBS with the current of 126.57*pA*. (C) The same as A, during IDBS. The black vertical thick lines indicate 50 mV.

#### EDBS and IDBS effects

Applying the EDBS (see materials and methods) to the STN neurons in the network model quenches the PD-like *β* oscillations in the STN (and in the other regions included in the network model; Figures 4A-C, top panels).

To quantify the performance of EDBS on the PD-like *β* oscillations in the network model, we measured the Fano factor and the oscillation index of the STN, GPe, and GPi population firing rates while the STN was stimulated. For EDBS, both Fano factor and oscillation index dramatically decreased compared to the PD state (*P* < 0.001, independent two-tailed t-test), for all regions measured (i.e., STN, GPe, and GPi; Figures 4E, F). We also compared the performance of the EDBS on improving the thalamic fidelity in the network model. We found that EDBS increased the thalamic fidelity (almost to the healthy level; Figure 4G). In addition, we found that the burst rate of the GPi neurons significantly decreased (*P* < 0.001, independent two-tailed t-test) compared to the PD and also to the healthy (Figure 5A). It matches the previous experimental^54^ and computational studies^61^ and indicates the validity of our model.

All in all, this result indicates that applying regular/irregular EDBS to the STN neurons in the network model, quenches the PD-like *β* oscillations and improves the thalamic fidelity to the cortical motor command.

Next, we tested whether high-frequency inhibition of the STN neurons (IDBS) has the same effects as the EDBS in the network model. To do that, we stimulated the STN neurons in the pathological network model by high-frequency inhibitory pulses (see materials and methods) and measured the PSD, Fano factor, and oscillation index of the STN population firing rate as well as the thalamic fidelity in the network model (Figure 4). Our simulation results show that IDBS (unlike the EDBS) fail to quench PD-like *β* oscillations and to improve the thalamic fidelity (Figure 4). Although the burst rate of the GPi neurons in regular and irregular IDBS slightly decreases (*p* < 0.001, independent two-tailed t-test) compared to the PD state, the GPi burst rate were still above the healthy state when the network model exposed to IDBS (Figure 5A). However, comparing the performance of EDBS and IDBS in the network model reveals that EDBS outperforms IDBS in improving PD signs (i.e., quenching PD-like *β* oscillations and improving the thalamic fidelity) in the network model. The question may arise whether the irregular IDBS (see Materials and methods) may quench the PD signs in the network model. To answer the question, we also applied the irregular manner of both DBS types to the STN (Supplementary figure 4). The results were the same as the regular manner of the DBS.

In the following, we try to explain why IDBS fails to improve the PD signs in the network model.

#### IDBS can not treat the pathological STN rebound bursting activity in the network model

We showed that only EDBS (and not IDBS) is able to quench the PD-like *β* oscillations in the network model (Figure 4). The reason why IDBS fails to quench the PD-like *β* oscillations in the network model is the rebound bursting activity of the STN neurons due to IDBS. In line with the experimental studies^88, 110, 111^, the STN neurons in the network model show rebound bursts of spiking activity after the strong inhibition. As the IDBS is a barrage of inhibitory inputs to the STN neurons in the network model (see Materials and methods), the STN neurons react to it by rebound bursts of spiking activities (Figure 6C). STN rebound bursts lead to an increase in the GPe spiking activity through subthalamo-pallidal excitatory pathway. Then, the higher GPe activity gives rise to inhibition of the STN neurons which results rebound bursting due to the T-type calcium current (Figure 6C). Such a mechanism retains the PD-like *β* oscillations in the network model during IDBS. Therefore, the reason why IDBS fails to quench the PD-like *β* oscillations in the network model is the propagation of the STN rebound bursts through the subthalamo-pallidal loop.

During EDBS, two behaviours can occur depending on the current amplitude. First, for 147.36*pA*, the STN neurons in the network model do not show rebound bursting activity (Figures 4, and 6A). The reason is that the STN spiking activity is driven by the EDBS pulses, counteracts with the pallido-subthalamic inhibition (see also Figure 2I), thereby no STN rebound bursting activity can happen due to the non-sufficient T-type calcium current. Second, for 126.57*pA*, the EDBS extends the burst durations and the inter-burst intervals which results quenching the PD-like *β* oscillations (Figure 6B).

### Excessive thalamic activity in resting state

#### Excessive thalamic activity occurs only in PD

So far, using our simulation results, we showed that the STN EDBS (and not IDBS) can quench pathological *β* oscillations emerging in the PD state of the network model. Next, we investigated the effect(s) of each stimulation type (i.e., EDBS or IDBS) on the resting state activity of the healthy and the PD network model. To simulate the resting state of the network model, we excluded the cortico-thalamic sensorimotor pulses (i.e., by setting the *I*_*SMC*_ in the equation 13 to zero for both healthy and PD network model simulations; see materials and methods). Our simulation results indicate that the thalamic neurons in the healthy network model, do not show any spiking activity during the resting state (Figure 7A). However, in the PD network model, the thalamic neurons show the mean spiking activity of 8.24±1*Hz* during the resting state (Figure 8C). This result seems to be reminiscent of the resting state tremor activity in the network model because of three reasons. First, such thalamic activity is specific to the PD state of the network model (compare Figures 7A and 7C). Second, according to the experimental studies^118–120^, the tremor is driven by the pathological thalamic activity in PD. Third, the resting state pathological thalamic activity in the network model approximately is 8 Hz which matches the tremor frequency reported in the experimental studies^65, 66, 89, 121–124^. However, our results do not replicate the thalamic tremor cell rhythmic activity similar to those experimental studies during tremor^65, 66^. Therefore, hypothesize that the thalamic excessive activity can be related to tremor but it does not represent the activity of the tremor cells. In the following, we explain why such thalamic spiking activity emerges only in the PD network model (and not in the healthy network model).

**Figure 7.**
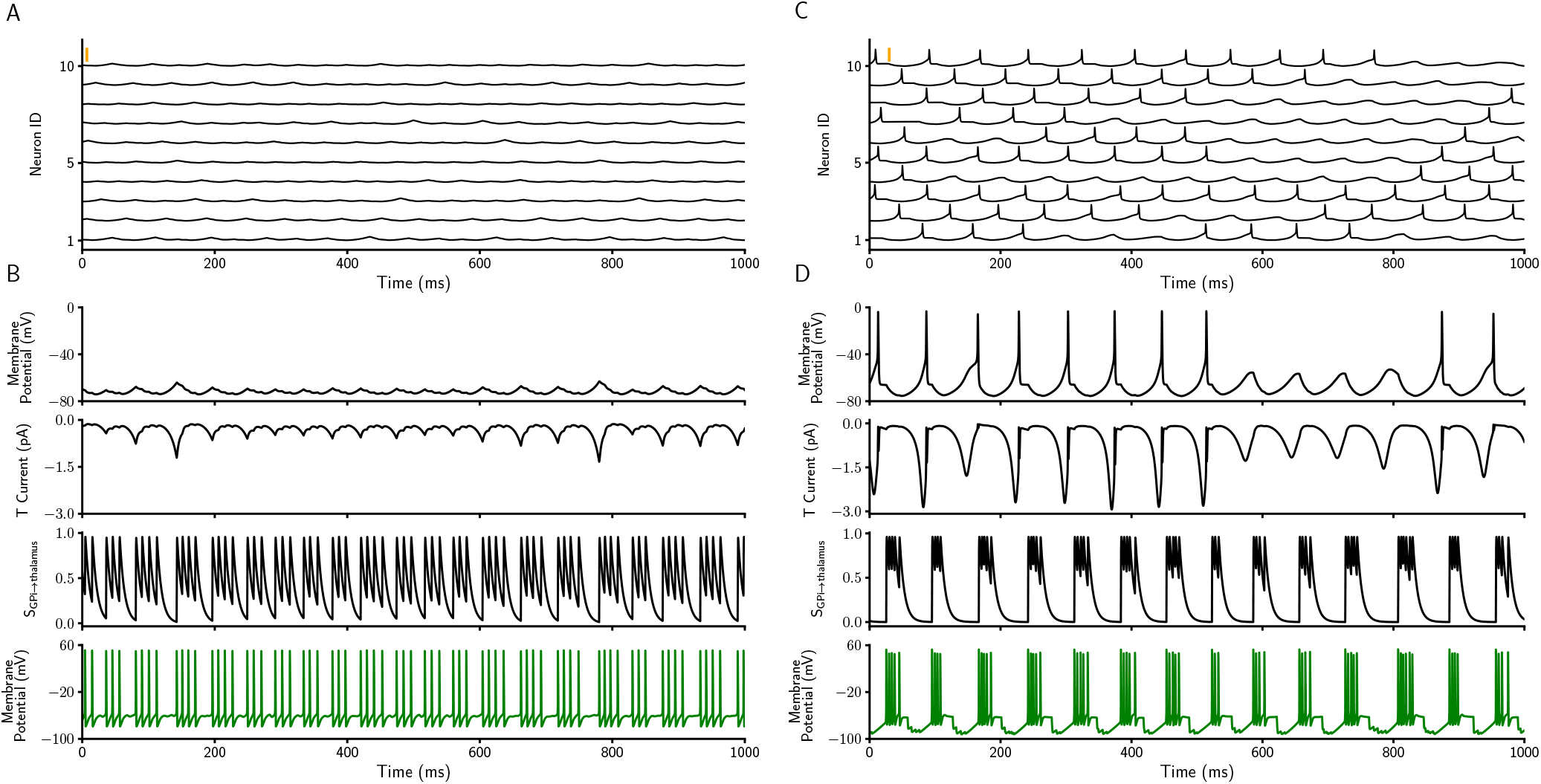
Excessive thalamic activity profile in healthy and PD state. (A) Membrane potential of 10 thalamic neurons in the healthy network model during resting state. The orange vertical thick line indicates 50 mV. (B) From top to bottom, membrane potential, low threshold T-type calcium current, synaptic inhibitory input from the connected GPi and membrane potential of the corresponding GPi neuron (see Materials and methods) in the healthy state of the network model. (C, D) The same as A and B for PD state of the network model.

**Figure 8.**
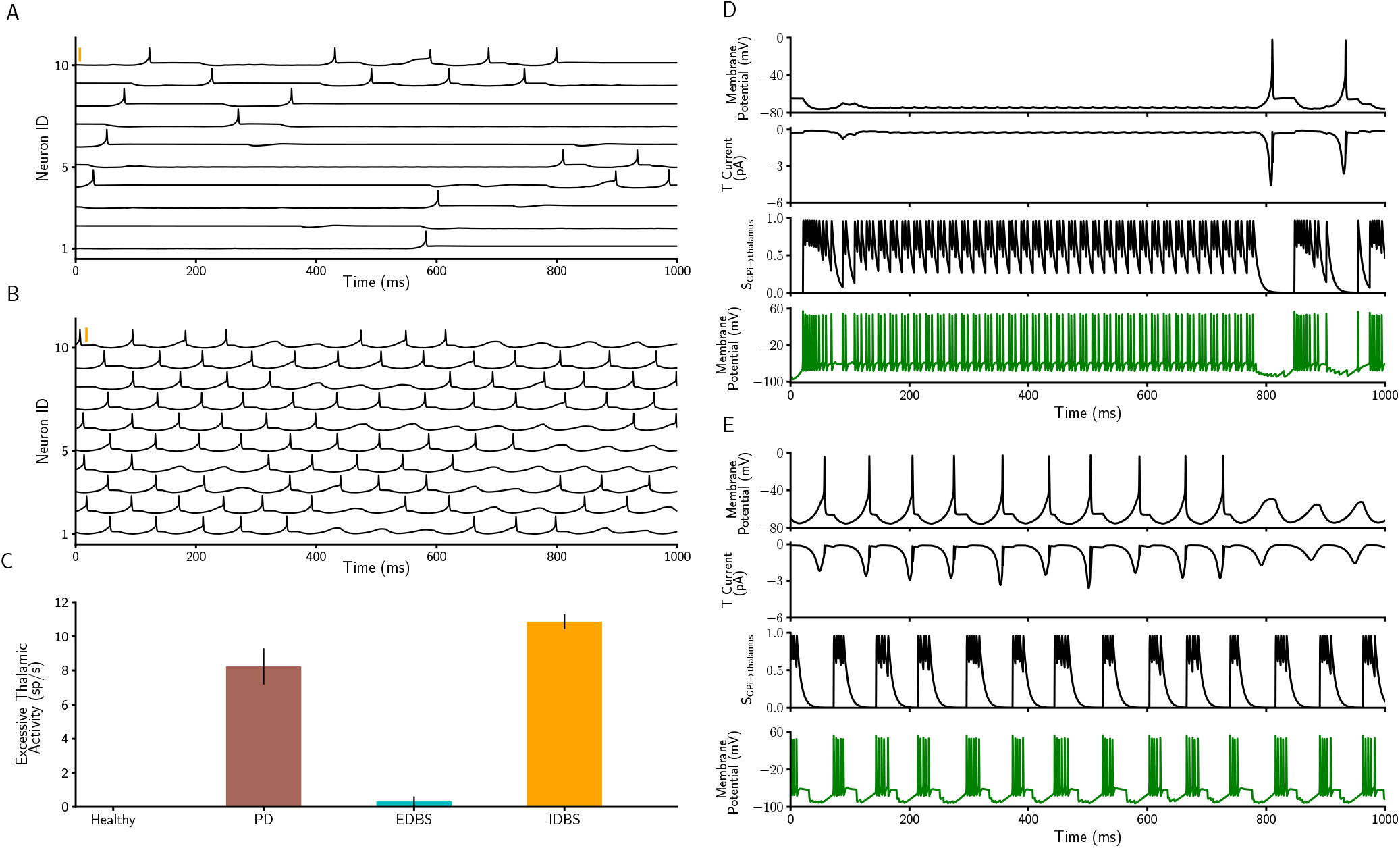
Excessive thalamic activity profile of PD state during EDBS and IDBS. (A, B) The membrane potential of 10 thalamic neurons when STN is exposed to regular EDBS (A) and regular IDBS (B) during the resting state of the network model. The orange vertical thick lines indicate 50 mV. (C) Excessive thalamic spike rate in the healthy and PD states and when the STN is exposed to EDBS and IDBS (error bars show standard deviation). (D) From top to bottom, membrane potential, low threshold T-type calcium current, synaptic inhibitory input from the connected GPi and membrane potential of the corresponding GPi neuron (see Materials and methods) when STN is exposed to EDBS in the network model. (E) The same as D, when STN is exposed to the IDBS.

The thalamic neurons in the network model receive inhibitory inputs only from the GPi neurons (Figure 1A; note that the cortico-thalamic input is set to zero during the resting state). Thereby, the resting state thalamic activity in the network model is driven by inhibitory inputs from the GPi. Our resting state simulation results show that the healthy GPi spiking activity is tonic (Figure 7B, bottom panel). Such tonic GPi spiking activity or weak bursting activity (burst with the lower number of spikes; Figure 7B, two bottom panels; and also see Figure5B and C) is not strong enough to sufficiently increase the T-type calcium current of the thalamic neurons. This leads to no thalamic spiking activity during the resting state and in the healthy condition of the network model (Figure 7B, top panel).

#### STN EDBS improves the excessive thalamic activity in the network model

Next, we investigated the effect(s) of EDBS and IDBS on the excessive thalamic activity in the network model. Mainly, we tested which DBS type (i.e., EDBS or IDBS) can reduce the resting state excessive thalamic spiking activity in the PD network model. Our simulation results show that applying EDBS to the STN neurons in the network model leads to better performance as compared to applying the IDBS (Figure 8). In other words, STN EDBS reduces the excessive thalamic spiking activity nearly to the healthy state while STN IDBS fails to do so (Figure 8A-C). In the following, we explain why STN EDBS outperforms STN IDBS in improving the excessive thalamic activity in the network model.

In line with previous experimental and computational studies^54, 61^, GPi neurons in the PD state of our network model switch between long-lasting hyperactivity and no activity states when the STN is stimulated by the EDBS (Figure 8D, bottom panel and also see the burst duration and number of spikes per burst in Figure 5B and C). During the GPi hyperactivity periods, thalamic neurons are strongly inhibited and thereby do not show excessive spiking activity anymore (Figure 8D). However, a single thalamic spike occurs at the end of the GPi activity (Figure 8D, top panel; also see Figure 8A). This single thalamic spike is due to the after-hyperpolarization increase in the T-type calcium current of the thalamic neuron receiving the GPi strong inhibitory input (Figure 8D).

However, when the STN is exposed to the IDBS, the GPi neurons in the network model show regular oscillatory bursts of spiking activity in (Figure 8E, bottom panel). These strong GPi bursts (similar property to PD state; see Figure 5A) cause sufficient flow of the T-type calcium current of the thalamic neurons in the network model (see the T current fluctuations in Figure 8E) which consequently, leads to excessive thalamic spiking activity. For answering the question whether the irregular IDBS improves the excessive thalamic activity, we investigated the thalamic activity when the STN exposed to both types of DBS with irregular pulses and the results show the same effect as the regular pulses (Supplementary figure 4G).

Altogether, our simulation results indicate that the EDBS outperforms the IDBS, not only in quenching the PD-like *β* oscillations in the network model (Figure 4), also in reducing the resting state excessive spiking activity of the thalamic neurons nearly to the healthy state (Figure 8C).

## Discussion

Permanent and excessive *β* oscillations in BG is the hallmark of the PD^11–13^. The DBS quenches *β* oscillations and improves motor symptoms related to PD^27–33, 36, 76, 112–117^. But, the mechanism of the DBS is poorly understood. In the present study, we investigated the generation of the pathological *β* oscillations in the BG by neural modelling and the effects of the DBS in different scenarios. We showed that the subthalamo-pallidal circuit is the potential source of the generation of the *β* oscillations based on the computational network model. Our findings confirm the rebound burst activity of the STN neurons is the key reason for to generation of the *β* oscillations as what has been suggested in^17, 23^. Then we investigated the role of the DBS in quenching the *β* oscillations. We applied inhibitory and excitatory DBS on STN neurons to compare their effect with those observed in experimental studies. Thereby, we tuned the current amplitude of each DBS types to have equal STN firing rates.

The results show that the EDBS can counteract the pallido-subthalamic inhibition and therefore, the rebound burst of the STN neurons was controlled (Figure 6). Also, EDBS can extend the duration of STN bursts which results in quenching the *β* oscillations (Figure 6). While the IDBS is not able to quench the *β* oscillations due to its ineffectiveness in the suppression of rebound burst activity of STN neurons and the extension of burst duration. Our results suggest that the effect of DBS on STN neurons is excitatory. Our simulation results also help to clarify the relationship between tremor and the activity of the BG and thalamus. We showed that the postinhibitory rebound activity of the thalamic neurons due to the strong inhibition of the GPi neurons is the potential reason for the resting state tremor. Like other signs of PD, the tremor activity of the thalamus was quenched by EDBS while the IDBS was worsened.

In the following, we discuss the network model, the generation of *β* oscillations, the tremor, and the role of DBS in improving the PD signs in detail.

### Network model

Our BG model has been created using the Hodgkin-Huxley type neuron model that can generate neural behaviour in detail (i.e., ion currents). Previous computational BG studies^17, 18, 57–62, 74, 125^ except^24, 126^ which were created by Hodgkin-Huxley type neurons, were not reported *β* oscillations in PD state. In these studies, one of the changes to set the model in PD state is strengthening subthalamo-pallidal synapses, based on the experimental results^69–72^. The PD *β* oscillations also were reported by the model proposed in^56^ which was created using the LIF neuron model. This model moves to the PD state only by increasing striato-pallidal spiking activity (without changing any synaptic strength). They show the PD *β* oscillations might arise from the network effects, without taking into account the details of neuronal dynamics. While ^23^ used integrate-and-fire-or-burst neuron in their network model for subthalamo-pallidal circuit. Their model moves to PD state by changes in synaptic strength and input spikes in subthalamo-pallidal circuit and represents the PD *β* oscillations. In this model, the rebound burst activity of the STN and GPe neurons played an important role in the generation of the *β* oscillations. The PD *β* oscillations have also reported in the firing rate models^20, 127–129^. These models cannot be used for single neuron study in the network. By contrast, the role of calcium bursting can not be demonstrated in these models. However, the modified BG model in the present study was able to generate the *β* oscillations in the PD state. Also, the other properties of the BG model (i.e., firing rates and changes in firing rates when moving from the healthy to the PD state) were matched the experimental observations. these modifications were made by changing some intrinsic and connectivity neuronal properties (see Materials and methods).

### Generation of beta oscillations

Rhythmic activity is a widespread dynamics of the brain circuits. While normal oscillations in healthy brain are crucial for transmission of the information between the brain regions^130–134^, abnormal synchrony disrupts efficient communication and is a hallmark of several neurological and cognitive disorders (epilepsy, PD, schizophrenia, etc). For example, enhanced *β* oscillations observed in the cortex and several nuclei of BG is a common trait of the brain dynamics in PD^6, 7, 9, 10^. However the source of these oscillations is still under debate. Computational and experimental studies have implicated the subthalamopallidal circuit as the potential source of the *β* oscillations in PD^7, 8, 14–24, 79^. Moreover, the induction of *β* oscillations from the cortex into the BG has been claimed by^6, 25, 26^. However, more detailed models for the generation of the *β* oscillations, the rebound burst activity of the STN neurons due to the inhibition of the GPe neurons in the BG has been considered^15, 16, 111^. Somehow, this hypothesis is confirmed when the motor symptom of patients with PD suppressed after receiving a T-type calcium channel blocker such as Zonisamide^135^. Also, the computational studies such as^17, 18, 23, 24, 126^ confirmed this hypothesis. While, the other studies demonstrate that the excessive inhibition on inhibitory population (such as striato-pallidal inhibition) and/or excessive excitation on excitatory population (such as cortico-subthalamic) result in *β* oscillations^20, 56, 74, 136, 137^. Our findings confirm the role of post-inhibitory rebound bursts of STN neurons in the generation of the *β* oscillations (Figure 2H and I).

### The role of the DBS is excitatory

The DBS improves PD-related motor symptoms^27–32, 115–117^ and PD-related neuronal behavior in BG^10–13, 33, 36, 76, 112–114^. In our network model, we quantified the PD-related signs by *β* oscillations, synchrony, thalamic fidelity, and tremor frequency (see below, materials and methods, and also Figure 2). Here, we applied two types of the DBS in our network model to see which satisfy our expectation about the amelioration of PD signs with DBS. Our findings suggest that the EDBS improved the PD neuronal behaviour and the motor symptoms while the IDBS worsened them (Figures 3 and 4). Only a small shift of *β* peak in PSD of the STN results from IDBS (Figure 4C). This shift may come from the change in burst rate and burst duration of STN due to extra inhibition despite the pallidal inhibition.

Although the previous studies^38–41, 43–47, 49, 50, 138^ indicate that the role of the DBS on its target is inhibitory, in^48, 51, 53–55^ has been demonstrated the opposite role by our findings. On the other hand, the computational study in^56^ has claimed the excitation of excitatory and/or inhibition of inhibitory populations leads the network to oscillations (in this case is *β* oscillations). Therefore, the pathological *β* oscillations have been quenched using high-frequency inhibitory spike trains on STN, which is not consistent with EDBS in our network. Still, there is opposite evidence for this study’s results: the initiation of movement accompanied with increasing cortical activity^139–142^ and the STN neurons are excited by cortex^105–107^ which quenches briefly the *β* oscillations in patients with PD^8, 143, 144^ and a computational model proposed in^24^. Moreover, the neuronal bursting activity was not investigated in that study, though in^145^ a computational model was proposed based on^56^ that the neuronal bursting activities have been investigated. This study showed the variety of STN burst range can affect *β* oscillations. In detail, the low burst rate of STN neurons quenched the *β* oscillations while the high STN burst rate generated the *β* oscillations with a little shift in frequency. Indeed, the intrinsic behaviour of STN neurons in a non-pathological state is bursty and the higher burst rate of BG nuclei in PD condition is also shown^146^ (and see also Figure 5 for GPi) which is consistent with the results reported in^145^. The consistency of this study and our network model is justified with considering that 1-the STN burst rate in our network model corresponds to high STN burst rate of the model in^145^ and 2-with considering the STN burst rate when it exposes to EDBS corresponds to low STN burst rate in that study. Consequently, it seems to the therapeutic effects of DBS acts by excitation of the STN.

However, in^147^ has been shown the GPe orchestrates the *β* rhythm in the BG network using the optogenetic method on mice. They showed either inhibition or excitation of STN does not decrease the pathological *β* oscillations in the network while the pallidal inhibition does. Also, they showed, the *β* induction in STN neurons using optogenetic excitation does not generate the *β* oscillations in cortical activity (ECoG), while the optogenetically induction of *β* in the GPe generates the *β* oscillations in ECoG. Inconsistency of these observations with our results may be rooted in the fact that continuous optogenetically excitation/inhibition may have different effects from those of high-frequency electrical pulses. Besides, the inhibition of the GPe results in the reduction of subthalamic inhibition which abolishes the STN post-inhibitory rebound bursts. Also, induction of the *β* rhythm in GPe by optogenetical excitation results in STN inhibition that causes STN rebound burst. These observations confirm the role of the STN rebound bursts in the generation of *β* oscillations in our study.

On the other hand, in^37, 148, 149^ it is shown that the DBS inhibits the STN neurons while it induces spike in their axons. In this hypothesis, despite the inhibition of STN neurons, their post-synaptic neurons (GPe and GPi) receive excitatory neurotransmitter (glutamate). So, inducing excitatory pulses in STN neurons in the network model matches this hypothesis. Our results suggested that the post inhibitory rebound burst of the STN neurons is the main cause of the *β* oscillations (Figure 2I). The EDBS by counteracting the pallidal inhibition in the network model eliminates the rebound bursting of the STN neurons, while the IDBS enhances the pallidal inhibition resulting in more rebound burst activity of the STN neurons in the network model (Figure 6). Altogether, based on our network model results we conclude that the role of the DBS on its target is excitatory (and not inhibitory).

The question may arise that the irregular manner of IDBS may quench the pathological *β* oscillations. Although a computational study has been shown the irregular DBS could not improve the PD signs^64^, we tested the effect of both EDBS and IDBS with irregular pulses and the results were approximately the same as regular pulses (Supplementary figure 4).

### Excessive thalamic activity in resting state

The tremor in patients with PD occurs when they are at rest. The frequency of the resting tremor in PD state is reported 3 to 8 Hz^66, 89, 121–124^ and it correlates with thalamic neuronal activity^118–120^. Previous computational studies^17, 18, 57^ demonstrated tremor activity by single neuron spiking activity in the tremor frequency in BG populations such as STN. While in^89^ the correlation between the high-frequency activity of STN and tremor has been shown. In a computational study, the tremor activity in the PD state has been represented by synaptic input from the GPi to the thalamic neurons in tremor frequency^62^. However, the resting state was not simulated in mentioned computational studies. In our network model, we simulated the resting state with interrupting sensorimotor command pulses to the thalamic neurons. Our results, show the extra thalamic spikes in tremor frequency in the PD state (and not in the healthy state).

In previous experimental studies it is shown that the rhythmic bursting activity of some of the thalamic cell is correlated with tremor^65, 66^. They called these thalamic neurons the tremor cells. The excessive thalamic activity in our network model did not match the rhythmic bursting activity with tremor cell. Moreover, with applying DBS in our network model the frequency of the excessive thalamic activity were decreased, while the previous behavioral studies have reported that by applying STN DBS the frequency of limb activity increased and while the amplitude of the limb movement was suppressed^78^. Therefore we hypothesize that the thalamic neurons in our model do not represent the thalamic tremor cells. Meanwhile, as the frequency of resting state excessive thalamic activities in our network model matches the tremor frequency and occur only in PD state, they might be related to tremor.

The resting state excessive thalamic activities were quenched when the network model exposed to the EDBS, while these activities were increased when the network model exposed to the IDBS. This finding, again, confirmed that the role of the DBS is excitatory.

## Conclusion

We utilized a computational model of the BG to investigate the underlying mechanisms of the DBS. With the help of the model we concluded that first, the rebound burst activity related to the T-type calcium current of the neurons has a key role in the generation of *β* oscillations. Second, we found that the role of the STN DBS is excitatory (and not inhibitory), because, the excitation of the STN neurons suppressed their rebound burst. Third, again, the rebound burst of thalamic neurons gave rise to the generation of resting tremor. Next, by exposing the STN neuron to high-frequency excitation the excessive thalamic activities-which may be related to tremor were quenched, while, exposing the STN neurons to high-frequency inhibition has been worsened. In summary, based on our model, we conclude that the role of the high-frequency DBS is excitatory on its target.

## Acknowledgements

We are grateful to Prof Dr Arvind Kumar and Dr Jyotika Bahuguna for their help and comments that improved our study. We are also grateful to Hamed Heidari, Davoud Nouri, and Dr Zeinab Fazlali for their helpful discussions. This work was partially supported by the Shahid Rajaee Teacher Training University.

## Supplemental Figures

**Supplementary figure 1.**
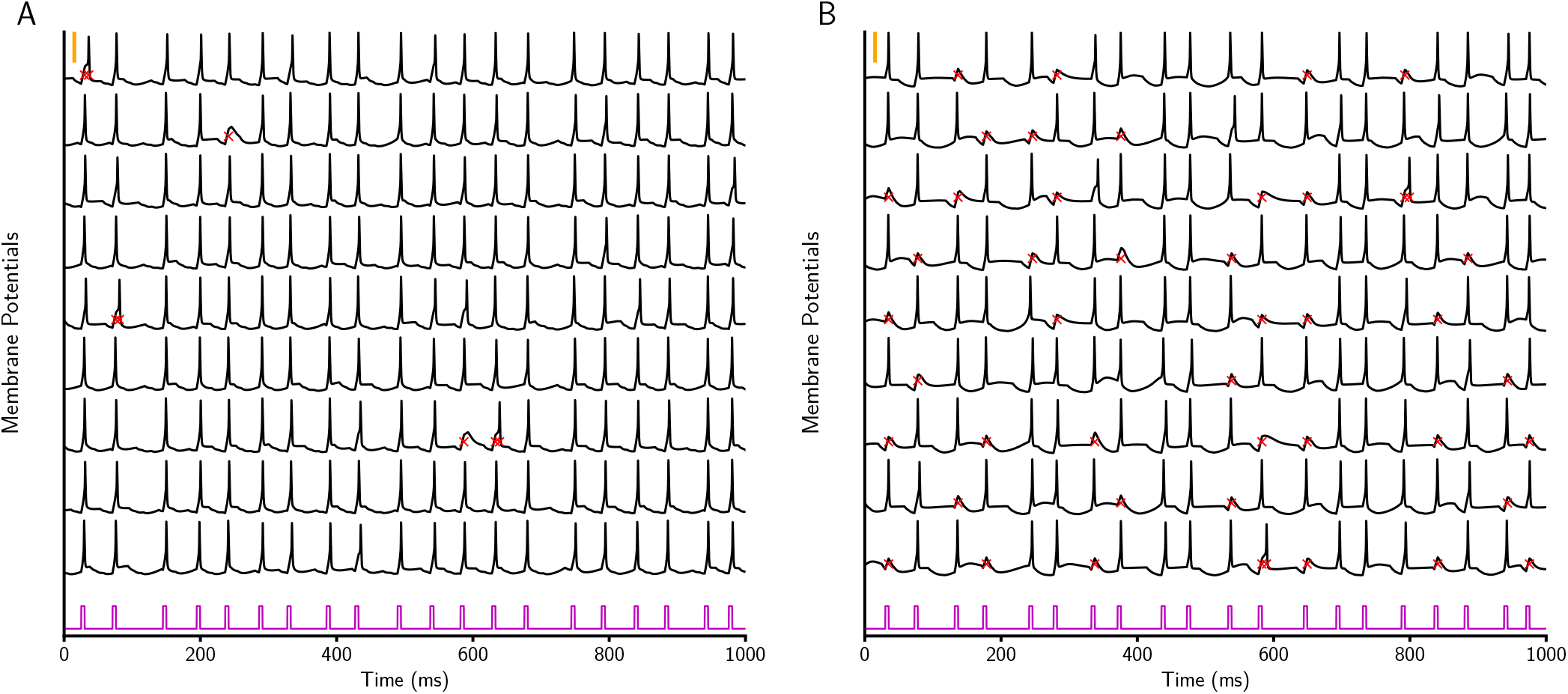
Thalamic response to cortical inputs. (A) Membrane potential of 10 thalamic neurons in the healthy state. The purple signal is the cortical input to each thalamic neuron. The red crosses indicate wrong or missed response to the cortical input. (B) The same as A for PD state. The orange vertical thick lines indicate 50 mV.

**Supplementary figure 2.**
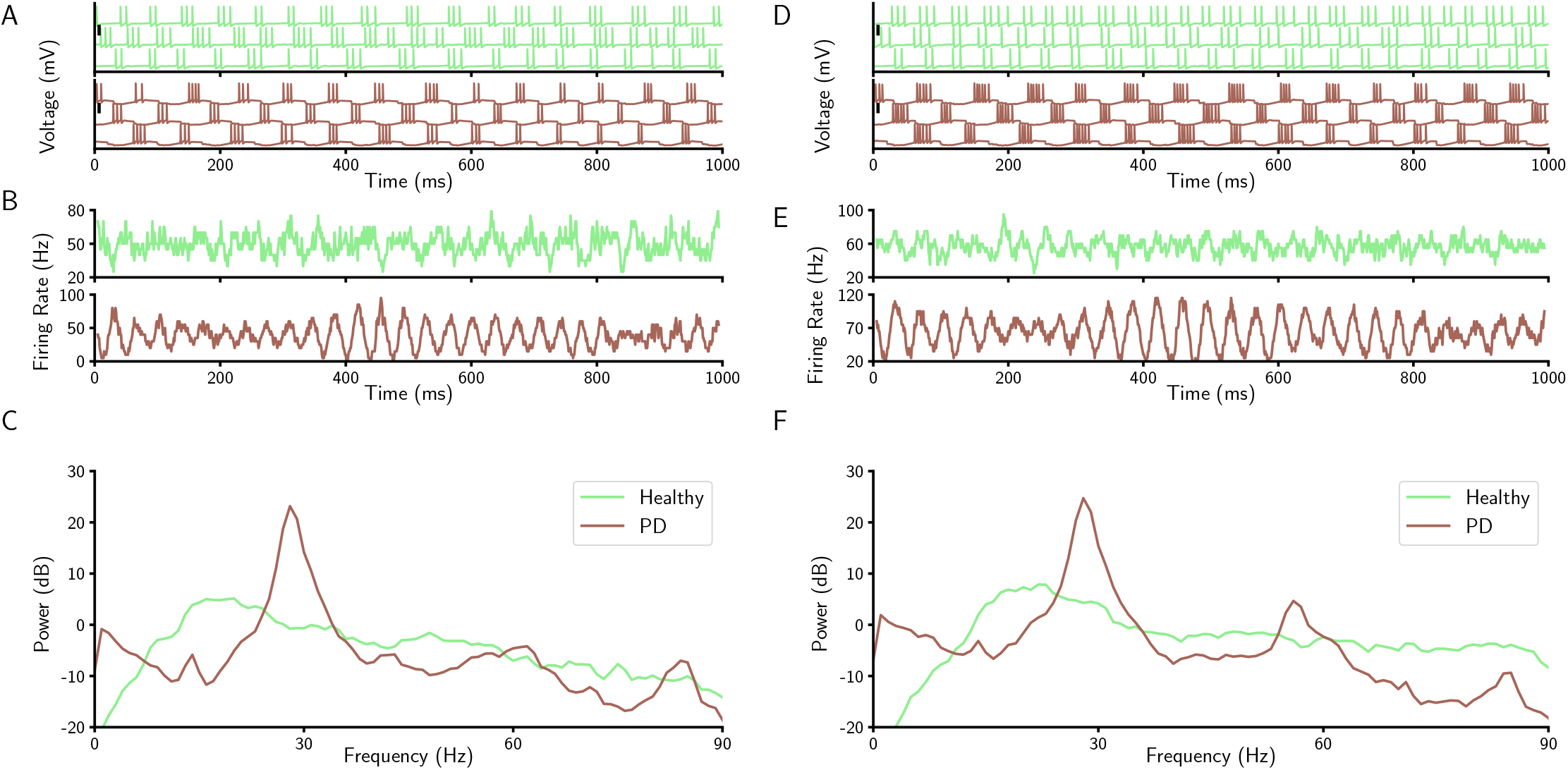
Neuronal and population properties of GPe and GPi in the healthy and PD states. (A) Membrane potential of three GPe neurons in the network model in the healthy (top) and PD (bottom). The black vertical thick lines indicate 50 mV. (B) Time resolved population firing rate of the GPe neurons in the healthy (top) and PD (bottom). (C) Mean power spectrum (average of 50 trials) of the GPe time resolved population firing rate in the healthy state (light green), PD state (brown). (D-F) The same as A-C for the GPi.

**Supplementary figure 3.**
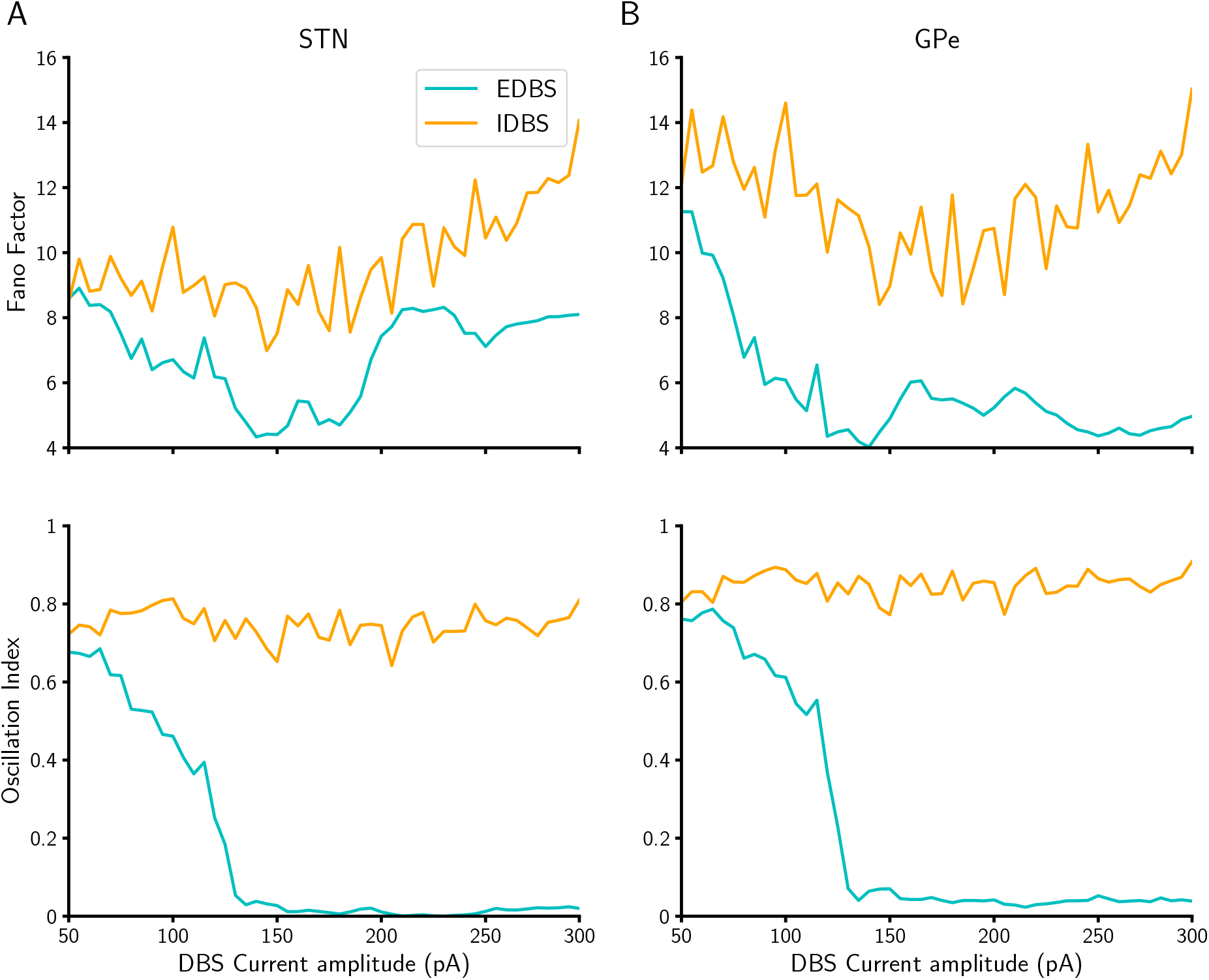
Fano factor and Oscillation index of STN and GPe across various DBS current amplitudes. (A) The fano factor (top) and oscillation index (bottom) of STN across various DBS currents amplitudes during EDBS (cyan) and IDBS (orange). (B) The same as A for GPe. Each point in all panels is averaged over 20 trials.

**Supplementary figure 4.**
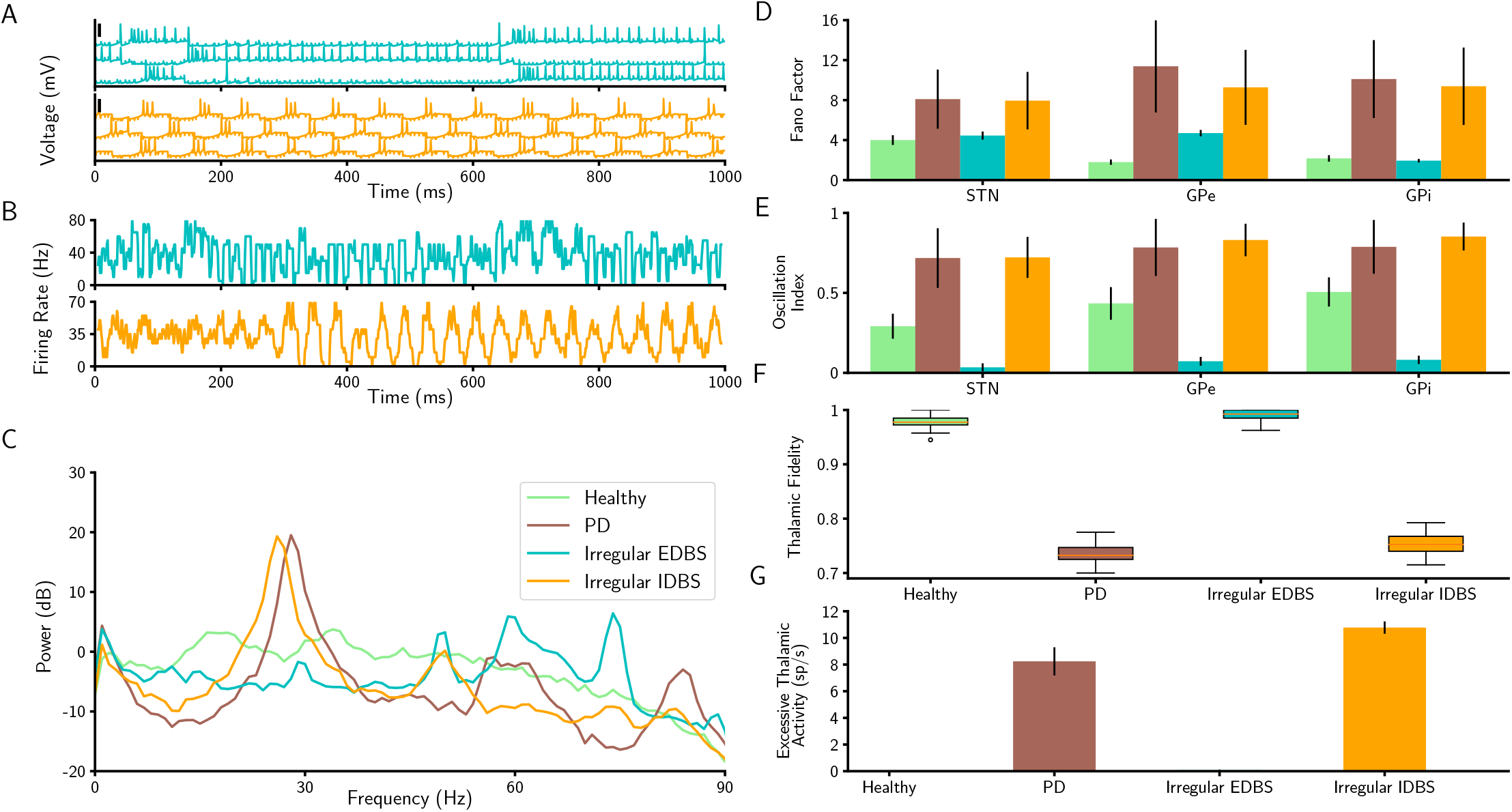
Neuronal and population properties of BG in Irregular EDBS and IDBS states. (A) Membrane potential of three STN neurons in the network model in the irregular EDBS (top) and irregular IDBS (bottom). The black vertical thick lines indicate 50 mV. (B) Time resolved population firing rate of the STN neurons in the irregular EDBS (top) and irregular IDBS (bottom). (C) Mean power spectrum (average of 50 trials) of the STN time resolved population firing rate in the healthy state (light green), PD state (brown), and during irregular EDBS (cyan) and irregular IDBS (orange). (D and E) Fano factor (D), and oscillation index (E) of the STN, GPe, and GPi in the network model (error bars show standard deviation; color codes correspond to C). (F) thalamic fidelity in the healthy state, PD state, and during regular and irregular IDBS. (G) Tremor-like frequency of thalamus in the healthy and PD states and when the STN is exposed to irregular EDBS and irregular IDBS (error bars show standard deviation).

